# Tonotopic and multisensory organization of the mouse dorsal inferior colliculus revealed by two-photon imaging

**DOI:** 10.1101/680603

**Authors:** Aaron B. Wong, J. Gerard G. Borst

## Abstract

The dorsal (DCIC) and lateral cortices (LCIC) of the inferior colliculus are major targets of the auditory and non-auditory cortical areas, suggesting a role in complex multimodal information processing. However, relatively little is known about their functional organization. We utilized *in vivo* two-photon Ca^2+^ imaging in awake mice expressing GCaMP6s in GABAergic or non-GABAergic neurons in the IC to investigate their spatial organization. We found different classes of temporal response, which we confirmed with simultaneous juxtacellular electrophysiology. Both GABAergic and non-GABAergic neurons showed spatial microheterogeneity in their temporal responses. In contrast, a robust reversed rostromedial-caudolateral gradient of frequency tuning was conserved between the two groups, and even among the subclasses. This, together with the existence of a subset of neurons sensitive to spontaneous movements, provides functional evidence for redefining the border between DCIC and LCIC.

## Introduction

The inferior colliculus (IC) is a major auditory processing center, which receives input from most brainstem auditory nuclei. The IC is usually divided into three main regions: the central nucleus (CNIC), the lateral cortex (LCIC), and the dorsal cortex (DCIC; Oliver 2005). The latter two, together also known as the “shell region” of the IC, are part of the non-lemniscal pathway, and are heavily innervated by feedback projections from the cerebral cortex. The DCIC is often defined as the part that covers the CNIC dorsally (e.g. Zhou and Shore 2006). The LCIC is characterized by distinct neurochemical modules, which can be visualized with various histochemical methods such as acetylcholinesterase or NADPH diaphorase staining as well as dense labelling of GABAergic terminals and sparse calretinin immunoreactivity (Lesicko et al. 2016; Dillingham et al. 2017). These modules show a distinct complement of Eph-ephrin guidance molecules during development (Wallace et al. 2016; Gay et al. 2018) and receive input from non-auditory areas (Lesicko et al. 2016; Patel et al. 2017), hinting at their integral role in multisensory integration in the IC. Here, we refer to the external cortex and lateral nucleus of the IC as the lateral cortex (LCIC; Loftus et al. 2008), which includes layer 3 or the ventrolateral nucleus, and to the dorsal cortex and pericentral nucleus as the dorsal cortex of the IC (DCIC). The DCIC and LCIC are at the surface of the mouse brain, making the dorsal IC an accessible structure for in vivo imaging (Geis and Borst 2013; Barnstedt et al. 2015; Babola et al. 2018).

Because of the prominence of descending, cortical input, the DCIC and LCIC is ideally studied in awake, behaving animals. A number of studies have addressed the firing of IC neurons in awake bats (e.g. Xie et al. 2005; Xie et al. 2007; Xie et al. 2008; Andoni and Pollak 2011; Gittelman and Li 2011; Gittelman and Pollak 2011; Gittelman et al. 2012) or mice (Gittelman et al. 2013; Grimsley et al. 2013; Duque and Malmierca 2015; Ayala et al. 2016; Grimsley et al. 2016; Longenecker and Galazyuk 2016; Galazyuk et al. 2017). However, the yield of such recordings is limited, and the acute nature of some of the studies means that residual anesthetics and analgesics may have interfered with neuronal activity.

Moreover, relatively little is known about different cell types in the DCIC and LCIC. Around one-fourth of neurons in the IC are GABAergic (Schofield and Beebe 2019). In particular the large GABAergic neurons seem to form a distinct subclass (Ito et al. 2009; Ito and Oliver 2012; Geis and Borst 2013), but based on histology, at least four different subclasses of GABAergic neurons can be discriminated, which all contribute to the ascending projections to the medial geniculate body of the thalamus (Beebe et al. 2018).

Here, we describe the use of two transgenic mouse lines to characterize GABAergic and glutamatergic neuronal subpopulations in the dorsal IC in awake animals using *in vivo* two-photon Ca^2+^ imaging. We studied GABAergic neurons with a Gad2-IRES-Cre mouse (Taniguchi et al. 2011) that was crossed with the GCaMP6s reporter line Ai96 (Madisen et al. 2015) and a sub-population of non-GABAergic neurons using the Thy1-driven GCaMP6s transgenic line GP4.3 (Dana et al. 2014). We show a rich diversity of sound-evoked responses in both GABAergic and non-GABAergic neurons in the awake mouse, confirmed with simultaneous juxtacellular electrophysiology in awake animals. Remarkably, we observed a reversal of the characteristic frequency gradient in the rostromedial-caudolateral direction, which was conserved between GABAergic and non-GABAergic cells, as well as among cells with different classes of sound-evoked response. Moreover, we found a subset of neurons that were responsive to spontaneous movement to the animal, and were potentially associated with multisensory neurochemical modules (Lesicko et al. 2016). These findings suggest that the large majority of the dorsal IC surface belongs to the LCIC.

## Results

### Expression of GCaMP6s in IC subpopulation with transgenic mice

To better understand the response heterogeneity observed in IC neurons, we attempted to divide the neuronal population through selective expression of the GCaMP reporter. Two transgenic mouse lines were used: GP4.3, a Thy1-driven GCaMP6s expression line (Dana et al. 2014), and a cross between Gad2-IRES-Cre line (Taniguchi et al. 2011) and Ai96, a Cre-dependent GCaMP6s reporter line (Madisen et al. 2015). From here on we will refer to the latter as Gad2;Ai96.

To confirm the neurochemical identity of GCaMP6s-positive neurons in the transgenic lines, we performed immunolabelling of GAD67. Figure 1 shows example staining and the proportion of GCaMP-positive neurons expressing GAD67 in each line. We have not distinguished between the different subregions of the IC in this analysis. The majority (4651/4898) of GCaMP-positive neurons in the GP4.3 line were not positive for GAD67 (Figure 1), showing that the Thy1-promoter selectively expressed GCaMP6s in non-GABAergic cells in the IC. Not all non-GABAergic cells were GCaMP-positive, as shown by the NeuN-positive, GCaMP-negative cell bodies in Figure 1C. In particular, we observed that GCaMP6s-positive neurons are enriched at the border of neurochemical modules characterized by dense GAD67 terminals (Figure 1D).

**Figure 1.**
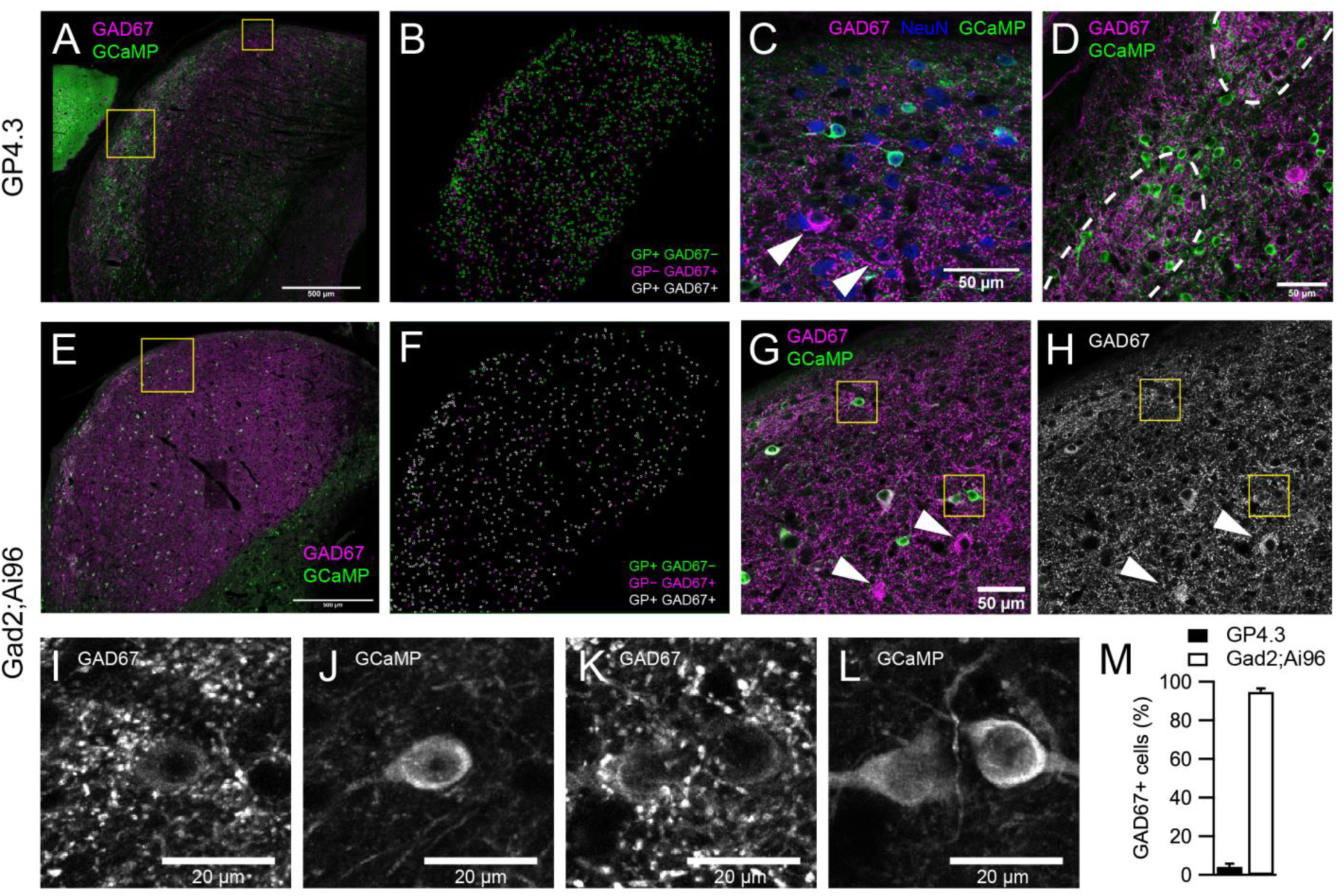
Co-expression between GCaMP6s and GAD67 in the two transgenic mouse lines. (A) Single optical section of an IC brain slice from a GP4.3 mouse immunolabeled for GAD67 (magenta) and GCaMP6s (green). (B) Distribution of GCaMP+ and GAD67+ cells in the 40 µm brain slice in (A), color-coded by their immunoreactivity to GFP and GAD67 antibodies. GP+/−: GCaMP-positive/negative. (C) Enlarged image from A (small square), showing different combinations of immunoreactivity: GCaMP (green), GAD67 (magenta) and NeuN (blue). Arrowheads point at two GP−GAD67+ cells. (D) Enlarged image from large square in A between two GAD67-dense modules marked by dashed lines. GP+ cells in GP4.3 were concentrated at the border of these modules. (E, F) Same as A and B, but from a Gad2;Ai96 mouse. (G-H) Enlarged region from E (square). Arrowheads point at two GP−GAD67+ cells. (I-L) Examples of GP+GAD67+ cells from G showing images from GAD67 & GCaMP channels separately. (M) Percentage of GCaMP+ cells showing immunoreactivity for GAD67 antibodies (error bars: s.e.m.; GP4.3: n = 3 slices from 3 animals; Gad2;Ai96: n = 4 slices from 3 animals).

In contrast, we detected positive GAD67 immunoreactivity in >93% (1702/1828) of GCaMP-positive neurons in the Gad2;Ai96 mice. Again, we observed that a fraction of the GAD67-positive cells in this mouse line were not labelled by GCaMP6s.

In addition to GAD67, the calcium buffer parvalbumin (PV) is another marker for neurochemical modules in the rat (Chernock et al. 2004), while the calcium buffer calretinin (CR) shows a complementary expression (i.e. in the extramodular region) in juvenile and early postnatal mice (Dillingham et al. 2017). We therefore investigated the co-expression of PV and CR in the two transgenic lines. We found that the large majority (87%) of GCaMP+ cells in GP4.3 were not positive for either marker (PV+: 4%; CR+: 8%; PV+CR+: 1%), whereas in the Gad2;Ai96 line a larger fraction of GCaMP+ cells were positive for CR or PV (PV+: 17%; CR+: 22%; PV+CR+: 7%; Figure 1 – figure supplement 1). At the single neuron level, the presence or absence of the two buffers was insufficient to indicate whether the neurons lay within a module.

### Sound-evoked changes in F_GCaMP_ of IC neurons

Unanaesthetized, head-fixed mice were subjected to 1-s pure tones of various frequencies and intensities (Figure 2). The use of longer stimuli allowed a detailed characterization of the kinetics of the fluorescence change (ΔF_GCaMP_). Importantly, we observed both increases, which were either transient or sustained, and decreases in F_GCaMP_ during the presentation of stimuli, and some cells showed a sharp increase in F_GCaMP_ directly after the cessation of the sound (Figure 2E). The different fluorescence response classes were classified as onset/sustained, inhibitory and offset, respectively (Figure 3A-D). The decrease in fluorescence upon sound stimulation was likely due to sound-evoked inhibition of spontaneous firing in these cells. Some cells showed a mixture of response kinetics, some even to the same stimulus. Particularly common was the combination of inhibitory and offset responses (e.g. Figure 3E,M). Another common combination were onset-offset cells in which a lower frequency elicited an onset/sustained response, while a slightly higher frequency elicited an offset response (e.g. Figure 3I). Inhibitory and offset responses were not reported in earlier imaging experiments performed in anesthetized mice (Ito et al. 2014; Barnstedt et al. 2015); their use of shorter stimuli did not allow a distinction between onset and sustained responses.

**Figure 2.**
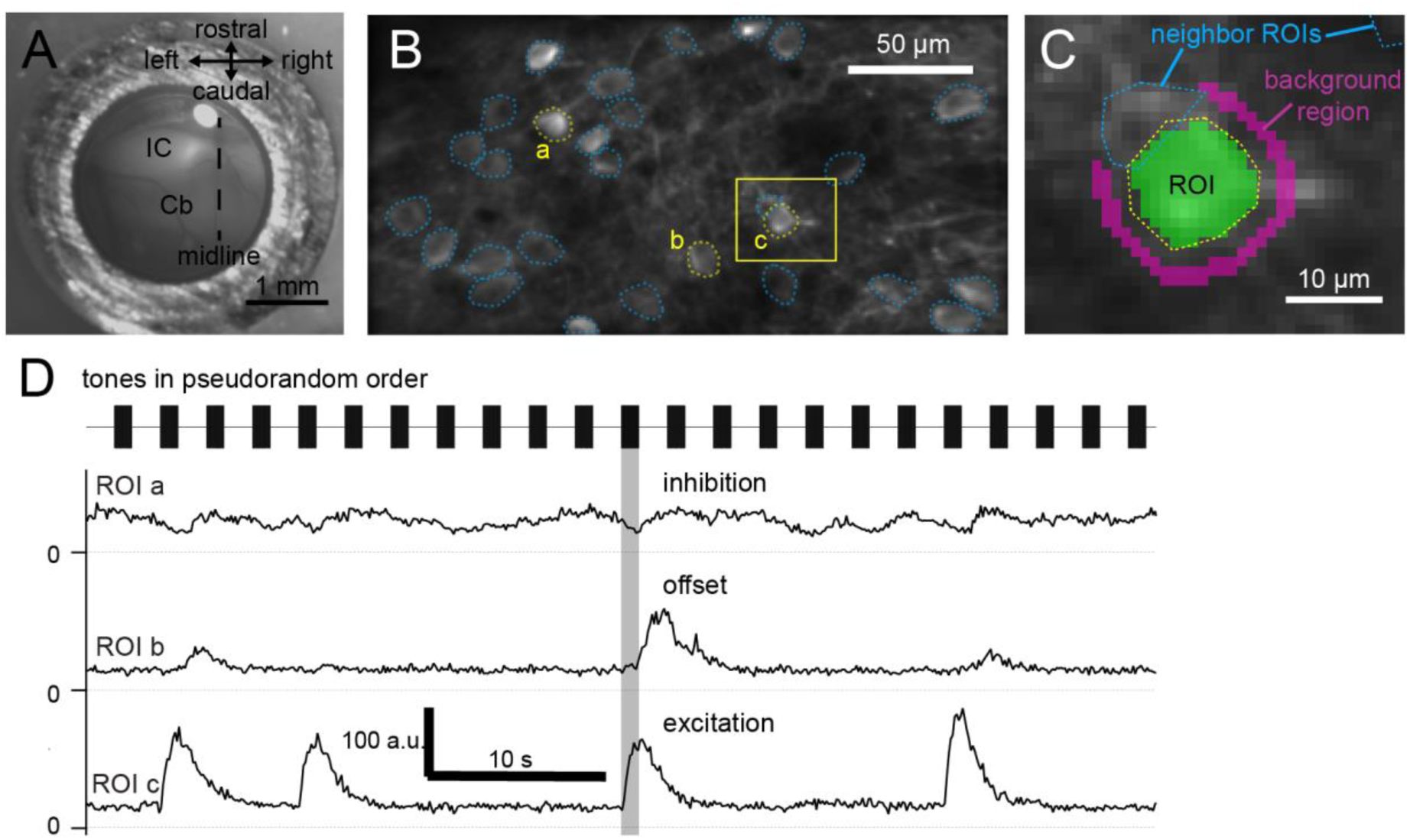
Calcium imaging of dorsal inferior colliculus in awake mice. (A) Top-down view of cranial window, showing optically exposed inferior colliculus (IC) and cerebellum (Cb). (B) Averaged GCaMP6s fluorescence in a 256 × 128 μm area, showing regions-of-interest (ROIs; dotted lines) defined around soma of neurons. (C) Enlarged view from C, showing pixels included in an ROI (green overlay) and a surrounding 2 μm wide background region (magenta overlay). Pixels belonging to other ROIs were excluded from the background region. (D) Background-subtracted fluorescence over time of the ROIs highlighted in C (a,b,c). Tones of 1-s duration with different frequencies and intensities were played, evoking different responses in IC neurons. The same sound (shaded area) evoked an inhibitory response in neuron a, an offset response in neuron b and a sustained excitatory response in neuron c.

**Figure 3.**
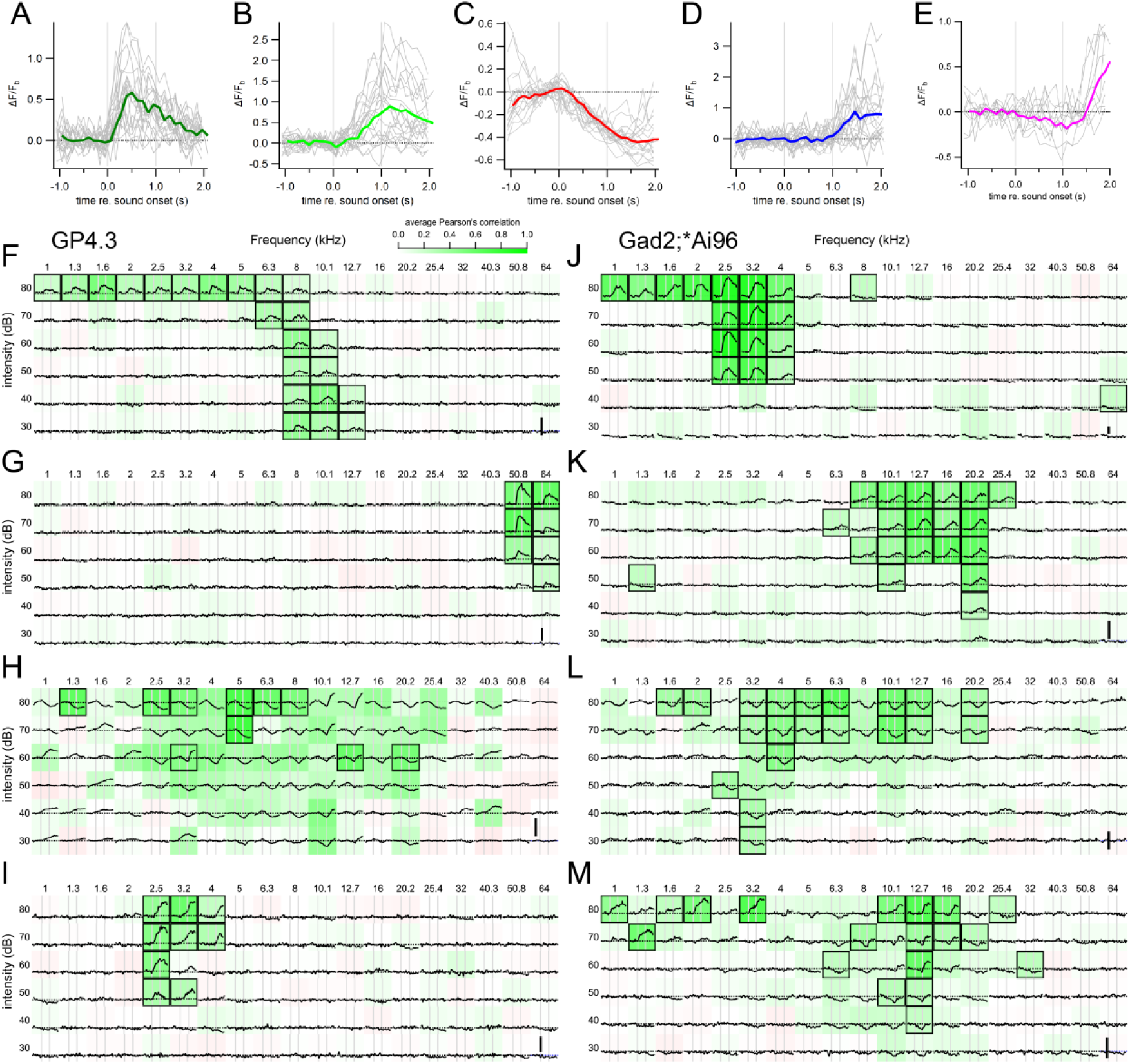
Different sound evoked responses and frequency response areas of representative example cells. (A) An onset fluorescence response to a 1-s pure tone. Grey vertical lines indicate onset and offset of sound stimulus. Colored and grey traces are average and individual trial fluorescence changes, respectively, normalized to a one-second baseline immediately before the stimulus onset (ΔF/Fb). (B) A sustained fluorescence response. (C) An inhibitory response. (D) An offset response. (E) A mixed inhibitory and offset response. Example frequency response areas from GP4.3 (F-I) and Gad2;Ai96 (J-M) mice. Each subplot shows the average ΔF/Fb to stimulus of the specified frequency and intensity. Background color shows average Pearson’s correlation among repetitions, indicating consistency of response (Geis et al. 2011). Black squares mark significant correlation from bootstrap analysis. Cells in (G) and (J) are examples of cells with onset fluorescence response; cells in (F) & (K) showed a sustained fluorescence response; cells in (H) & (L) were inhibited by sound; Cells in (I) and (M) shows a mixture of different response classes. (I) A typical onset-offset cell with frequency dependent responses: onset fluorescence responses to 2.5 kHz tones, and offset fluorescent responses to 3.2 and 4 kHz tones. (M) A cell showing intensity-dependent responses: at 12.7 kHz, low intensity tones evoked a decrease in fluorescence (inhibited) while tones at 80 dB elicited an offset response. Tones at 60-70 dB elicited a mixture of inhibition and offset response. Vertical scale bars in (F-I) indicates 1 Fb.

In GP4.3 mice, 585 out of 1017 cells showed detectable fluorescence changes to 1-s pure tone stimuli (see Methods). Among the responsive cells, 188 cells (32%) showed a pure onset/sustained response (e.g. Figure 3 – figure supplement 2), with fluorescence increasing only during sound presentation; while 64 cells (11%) showed a pure offset response, with fluorescence increasing after sound presentation. Another 253 cells (43%) showed a pure inhibitory response, with fluorescence decreasing during sound. The remaining 80 cells (14%) showed mixtures of the three response categories. The proportion of different response types are summarized in Figure 4A. The proportion of cells showing sound-evoked inhibition was clearly higher than the 2-23% reported for anesthetized C57BL/6 mice in Willott et al. (1988a).

**Figure 4.**
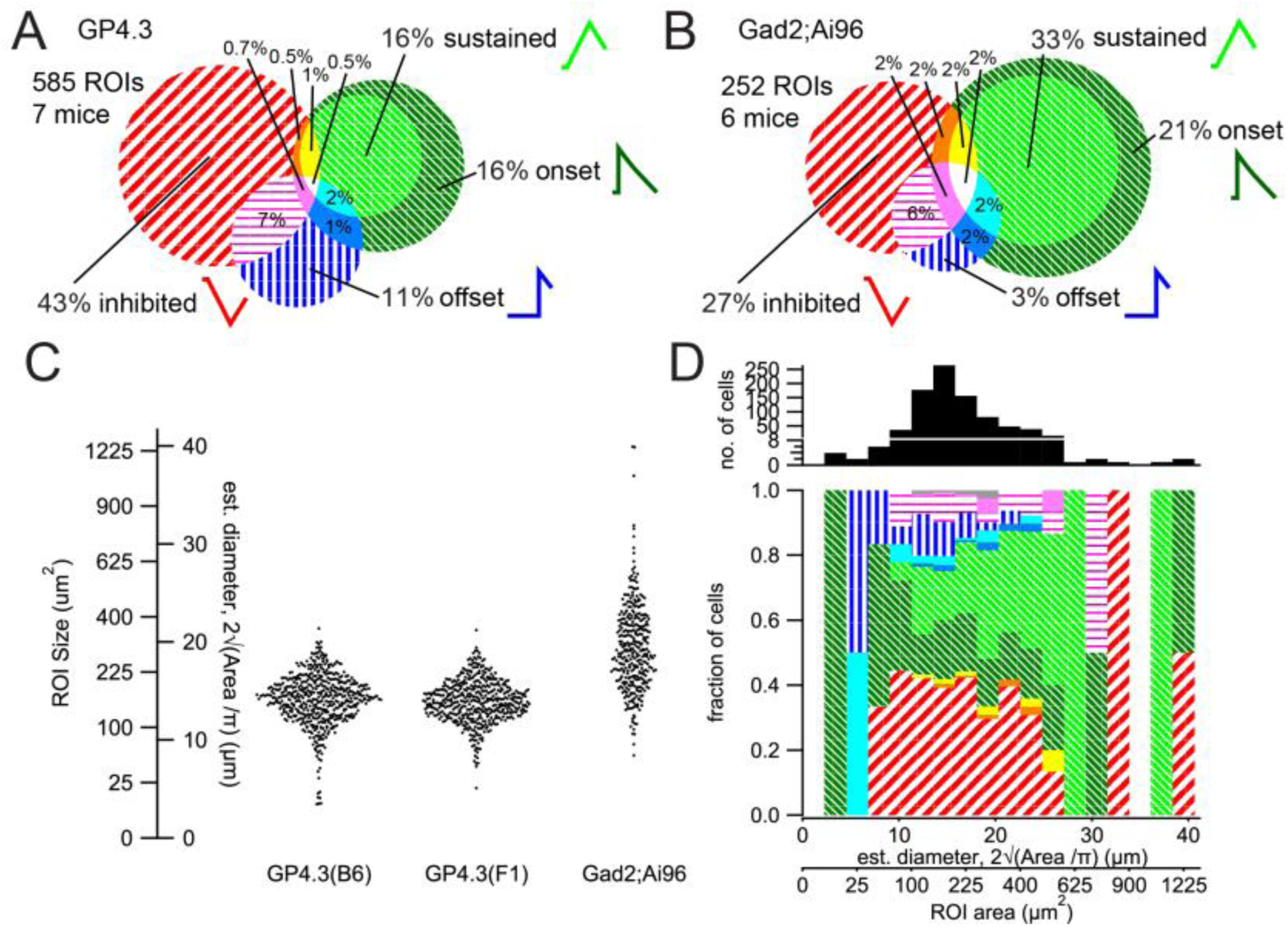
Proportion of response classes and relationship with cell size. (A-B) Proportions of response classes in GP4.3 (A) and GAD2;Ai96 animals (B). (C) Bee swarm plot for cell size for the different genotypes, measured as the size of the ROI (*A*) in imaging experiments. Estimated diameters (*d*) were calculated by assuming a circular shape (i.e. 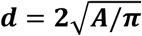). Cells imaged in Gad2;Ai96 mice were on average larger than those in GP4.3 mice. (D) Proportion of different response classes for cells of different sizes. There were more sustained responses in cells with a larger size.

Of a total of 342 GCaMP6s-positive IC cells in IC neurons from Gad2;Ai96 mice, 252 cells showed consistent fluorescence responses to 1-s pure tone stimuli. Among the responsive cells, 135 cells (54%) showed a pure onset/sustained response. Eight cells (3%) showed a pure offset response. Another 69 cells (27%) showed a pure inhibitory response. The remaining 40 cells (16%) showed mixtures of the three response categories. The proportion of different firing types in Gad2;Ai96 mice are summarized in Figure 4B.

### Relationship between cell type and soma size

GABAergic cells had a larger soma size (Figure 4C), defined as the area of the ROI in imaging experiments, as had been reported previously (cat: Oliver et al. 1994; rat: Merchán et al. 2005). In our dataset there was a tendency for larger cells to have a more sustained response (Figure 4D).

### Detailed kinetics of the different response classes

The temporal kinetics of the F_GCaMP_ response of each cell was assessed by averaging the fluorescence change across all stimuli that showed a significant response. Single exponential functions were fit on the onset (0-1 s re stimulus onset) and the offset/decay (0.5-4 s re stimulus offset) periods (Figure 3 – figure supplement 1A,B). Due to the limited stimulus duration, any fit resulting in an onset time constant (τ_onset_) greater than 1000 ms to an increase in fluorescence was considered sustained activity and these fit constants were not used for averaging. For excitatory responses (onset, sustained and offset), a time constant of around 1 s (Figure 3 – figure supplement 1D) was found for the decay period, similar to the decay time constant reported previously for GCaMP6s (Chen et al. 2013).

### Electrophysiological correlates of response classes

To relate the different fluorescence response to spiking patterns, which are highly heterogeneous within the IC (e.g. Willott et al. 1988a; b; Tan and Borst 2007), we combined Ca^2+^ imaging with *in vivo* juxtacellular recordings in awake mice. We recorded from neurons showing onset, sustained and inhibitory tone-evoked fluorescence responses. For fluorescence responses that we classified as onset, electrophysiological recordings showed that it can represent a cell that quickly adapted during our one-second long stimuli (example in Figure 5A-D). We observed sustained fluorescence responses that corresponded to sustained firing (Figure 5D) with different amount of adaptation. Inhibitory responses corresponded to cells that reduced their spontaneous firing upon sound stimulation (Figure 5D).

**Figure 5.**
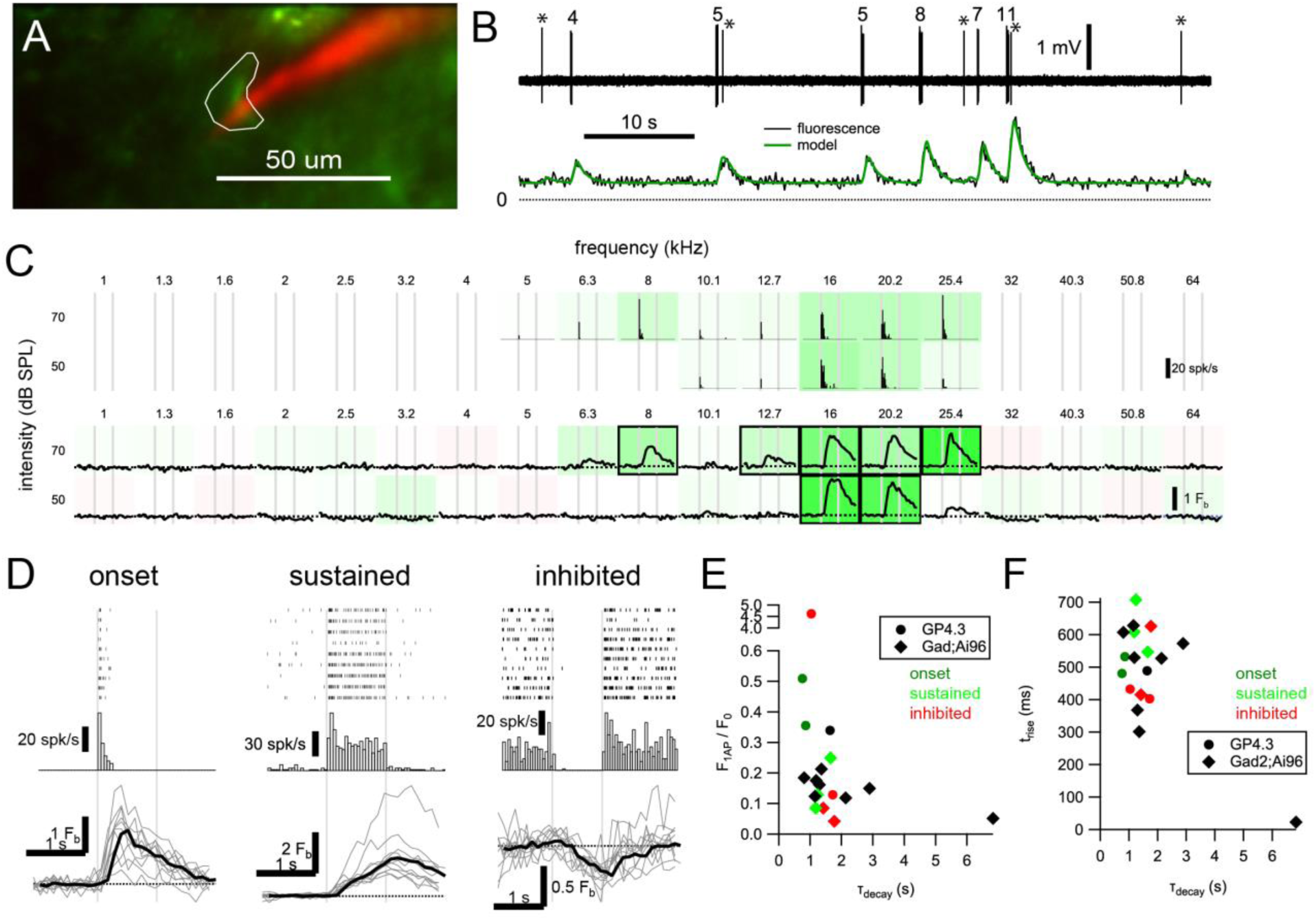
Relationship between GCaMP6s fluorescence and spikes in IC neurons. (A) Imaging of a GCaMP6s+ cell (green) in an awake GP4.3 animal with simultaneous juxtacellular recording with a pipette filled with Alexa 594 (red). (B) Spiking pattern (upper) and fluorescence (lower) of the example cell can be well related by a linear convolution of spike rate with a linear ramp, exponential decay kernel (model: green). Fit parameters: F0 = 28.8 a.u.; ΔF1AP = 10.2 a.u.; trise = 532 ms; τdecay = 859 ms. Numbers above bursts indicate number of spikes, and asterisks (*) mark single spikes. (C) Frequency tuning of the cell. Peristimulus time histogram (PSTH; upper, 50 ms bins, 10 repetitions) and fluorescence change (mean: black; individual: gray) of the cell in response to 1-s tone bursts. (D) Raster plot, PSTH and fluorescence response of an onset cell (same example as A-C), a sustained cell and an inhibited cell. Due to their rarity, we have not obtained a simultaneous recording for offset cells. (E) Relation between normalized single action potential amplitude (F1AP/F0) and decay time constant (τdecay). (F) Relation between rise time (zero-to-peak; trise) and τdecay. Color in E & F indicate response class (red: inhibited; dark green: onset; light green: sustained; black: no tone-evoked response) of the cell. Figure 6

We further characterized the relationship between fluorescence responses and firing patterns by fitting a model to our data set (Figure 5E,F; see Materials and Methods for equations). We found that on average, an action potential led to a median increase of 0.35 (GP4.3; n = 5 cells) or 0.14 (Gad2;Ai96; n = 12 cells) times the minimal fluorescence (F0), with a median zero-to-peak rise time of 480 ms (GP4.3) or 560 ms (Gad2;Ai96) and a median decay time constant of 1.04 s (GP4.3) or 1.36 s (Gad2;Ai96). The model explains between 50% and 95% of the variance in the fluorescence (median: 79%; average ± s.d.: 77 ± 13%).

### Spatial distribution of frequency tuning

The widespread and homogeneous expression of GCaMP6s in IC of the transgenic mice allowed a good overview of its functional organization. We aligned the position of the cells from multiple animals (GP4.3: n = 7 mice; Gad2;Ai96: n = 6 mice) by anatomical landmarks (midline, anterior and posterior extent of the exposed IC, lateral extent of the exposed IC), and plotted the cells on this common anatomical coordinate system (Figure 6A; Video 1).

**Figure 6.**
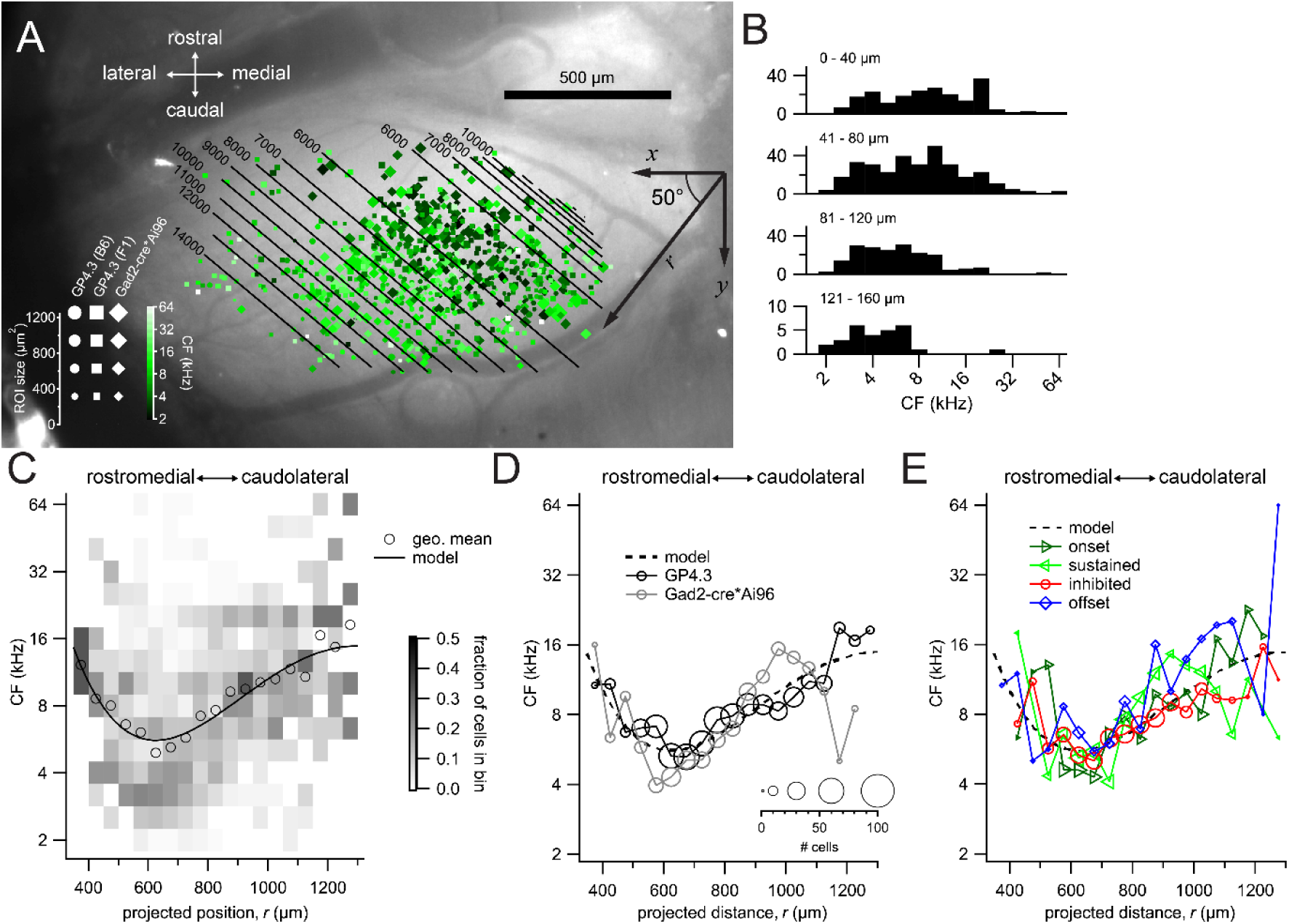
Tonotopic organization. (A) Combined spatial distribution of characteristic frequencies (CFs) in the two transgenic lines, aligned to the same top-down image of an exposed left IC. Symbol size represents the size of the ROI, while darker and lighter colors indicates lower and higher CFs, respectively. Cells from Gad2;Ai96 mice were marked by diamonds. For GP4.3, the shape of the symbol represents ROIs from C57BL/6J (circles) or B6CBAF1 (squares) background. To capture a direction of tonotopy, CF was fitted with a 4^th^-order polynomial 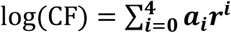 where *r* is a presumed direction of tonotopy at an angle *θ* (i.e. ***r* = *xcos θ + ysin θ***), with the origin being the contact point between superior and inferior colliculi at the midline. The best fit yielded an angle *θ* of 50° from the medial-lateral axis. Contour lines indicate the predicted CF of the fit. (B) Histogram of characteristic frequency grouped according to depth from pia surface. Note that low CF cells are present at all imaging depths. (C) Geometric mean (open circles) and probability distribution (background shading) of CF along the 50° line, in 75 μm bins. Black trace shows the fitted 4^th^-order polynomial. (D) Geometric mean of CF along the 50° line for the GP4.3 (black) and Gad2-cre*Ai96 (gray) lines. Size of symbol represent number of cells in each 75 μm bin. (E) Geometric mean of CF for cells of pure onset, sustained, inhibited or offset response classes.

Interestingly, we observed a central strip running in the caudomedial-rostrolateral orientation of cells responding to lower frequency, while CF progressively increased in both caudolateral and rostromedial directions. We tried to estimate the orientation of the spatial organization by projecting the x,y-coordinates of all neurons onto an axis with a parametrized angle θ, while simultaneously fitting the log-transformed CF using a polynomial model (see Materials and Methods). We used a polynomial model as a generic fit as a means to extract any general direction along which CFs seem to diverge, without imposing any presumption about the spatial dependence of CF. We found that a fourth order polynomial captured the variance maximally; further increasing its order did not lead to better fits. The results of the fit are presented as contour lines on Figure 6A, with the best orientation (θ) 50° from the medial-lateral (x) axis.

Barnstedt et al. (2015) suggested from clustering analysis that these low frequency neurons were from the most dorsal end of the central nucleus. While we do not exclude the possibility of imaging into the central nucleus of the IC, we believe the observed tonotopy is a good representation of the IC shell region because all neurons reported in this data set did lie within 160 μm from the pia surface. In addition, all CFs were represented among the most superficial neurons (<40 μm deep, Figure 6B). There was, however, an over-representation of low CF neurons among the deepest imaged (121 – 160 μm deep, Figure 6B). We attribute this to a sampling bias where the low frequency region coincided with the center of the cranial window, thus providing the best optical access for deeper imaging.

To better visualize the reversal of tonotopy, we projected the neurons onto our best orientation (Figure 6C-E). There was a high count of low CF (∼4 kHz) neurons near the 600 μm position (Figure 6C), with the mode going towards ∼20 kHz at the 1300 μm end. The increase in CF on the rostromedial side was not as pronounced, likely due to the low number of neurons sampled in this region. The solid line in Figure 6C shows our best polynomial fit, which explained around 13% of the variance in CFs. A similar spatial distribution of CF was observed for GABAergic and glutamatergic neurons (Figure 6D) and for cells showing different response classes (Figure 6E).

### Spatial distribution for response classes

We next asked whether there was any obvious spatial organization of the different response classes. Figure 7A shows the proportion of different response classes along our presumed tonotopic axis, which looked rather homogenous, without any obvious difference between cells rostromedial and caudolateral to the CF minimum. The same pattern holds for the orthogonal direction (Figure 7B).

**Figure 7.**
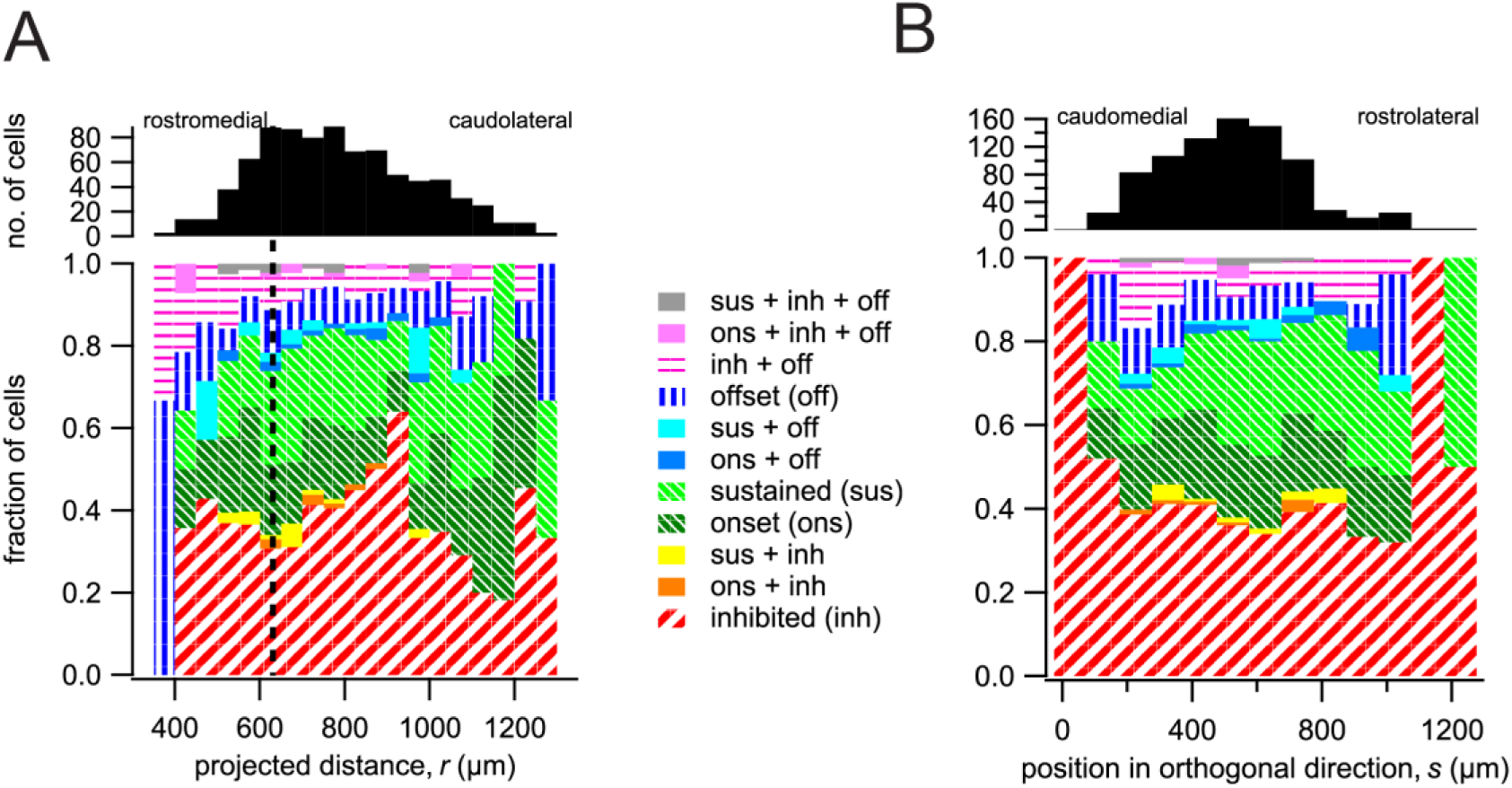
Spatial organization of response classes. Similar proportions of response classes along (A) the presumed tonotopic axis (bin size: 50 μm) and (B) the direction orthogonal to the axis (bin size: 100 μm). Zero position was taken as the contact point between superior and inferior colliculi at the midline. Dashed line in (A) denotes the minimal CF position based on our polynomial fit (625 μm).

### Neurons with movement related activity and their localization

Interestingly, we observed cells that showed fluorescence transients in the absence of sound presentation (Figure 8A). Numerous studies have shown both ascending and descending somatosensory projections to the IC (Aitkin et al. 1981; Künzle 1998; Jain and Shore 2006; Zhou and Shore 2006; Lesicko et al. 2016; Patel et al. 2017), as well as from nuclei that are upstream from IC such as the dorsal cochlear nucleus, which also receive somatosensory inputs (Wu et al. 2014). This prompted us to investigate whether these “spontaneous” activities could be attributed to non-auditory inputs. Our recordings were performed while the animal was passively awake, which allowed us to observe and correlate the voluntary movement of the animal to simultaneously recorded calcium transients.

**Figure 8.**
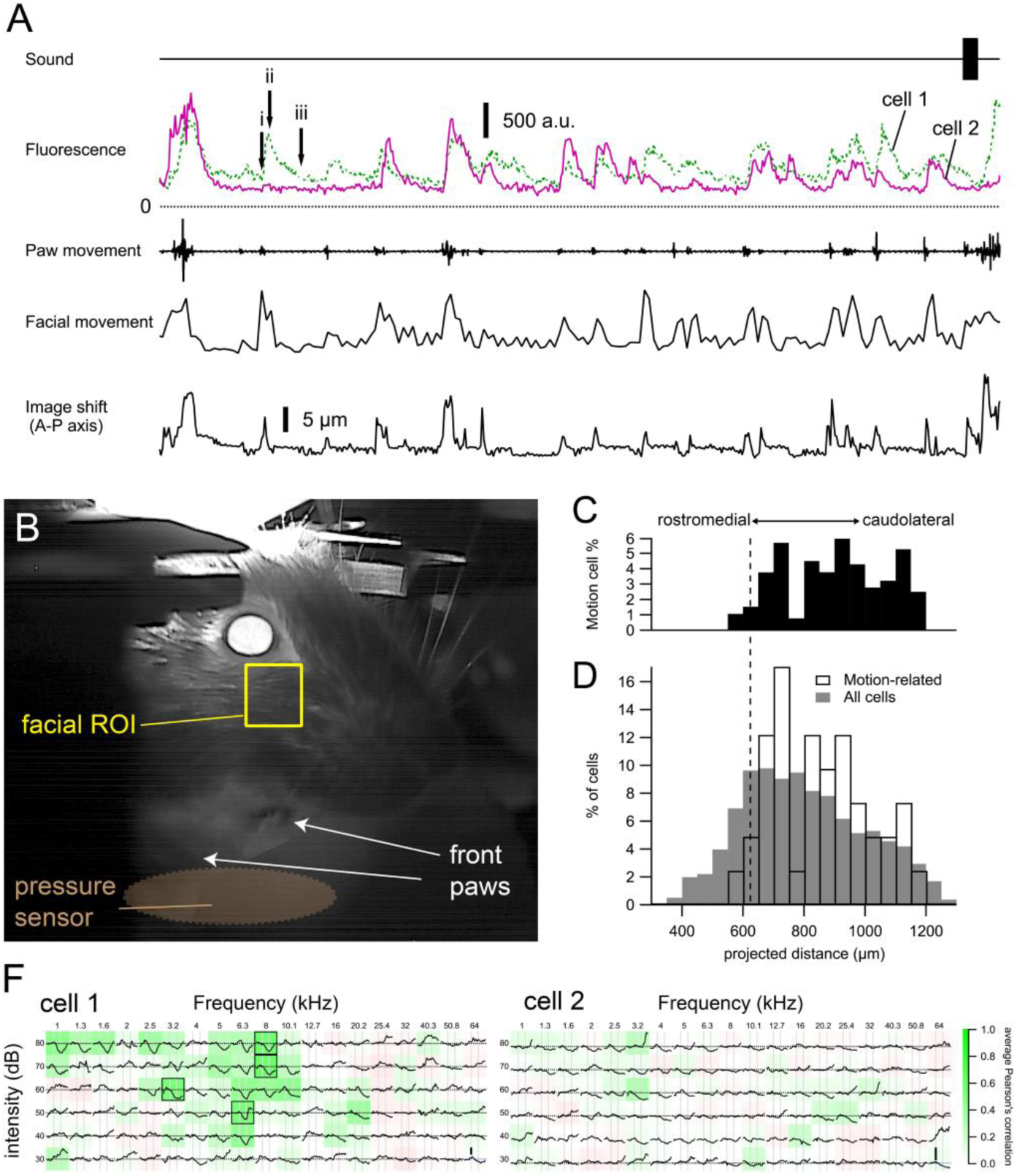
Cells with motion-related responses. (A) Two example cells that were spontaneously active in the absence of sound presentation. Calcium transients occurred when the animal was moving its paws or showing facial movement (e.g. whisking). These transients decayed much more slowly than motion artefacts (image shifts, bottom trace; see also Figure S3). (B) Facial movements were quantified from a simultaneously recorded video of the animal by calculating the root-mean-square of the changes in pixel intensity between consecutive frames of a rectangular area at the whisker pad. (C-D) Cells showing motion-related activity were predominently located in the caudolateral part of the dorsal IC. Broken line indicates the minimum CF position obtained from the polynomial fit. (F) The motion-related calcium transients were unlikely to be caused by sounds, as the two cells showed either an inhibitory FRA (cell 1) or no clear tone-evoked response (cell 2).

We found that the onset of many of these spontaneous transients coincided with movement events of the animal (paw and facial movements in Figure 8A). By correlating movement and fluorescence, we detected 165 (out of 1359) cells that showed a positive correlation (*r* > 0.25) with facial movements during the spontaneous recording period. To exclude potential false-positive detection due to motion artefacts (i.e. cells moving into and out of focus), we excluded cells with fluorescence transients that did not show the typical ∼1 s exponential decay kinetics, which last much longer than the brief image shifts that could accompany movement events (image shifts in Figure 8A, Figure 8 – figure supplement 1). Since animal movement inevitably produces sound that may activate the IC neurons, leading to an apparent movement sensitivity, we also excluded the cells that showed excitatory or offset responses in their FRA. In the end, 41 cells showed spontaneous calcium transients that correlated with animal movement and that could not be explained by their FRA, which was either inhibited or showed no clear pure tone evoked responses (example FRAs in Figure 8F). These cells seemed to be enriched at the caudolateral side of the dorsal IC (Figure 8C-D). Four of the 41 cells were found in the Gad2;Ai96 line, in agreement with a recent report that somatosensory inputs can directly target GABAergic cells (Olthof et al. 2019), and the remainder in the GP4.3 line, making them 1% of GABAergic and 4% of investigated glutamatergic cells, respectively.

### Comparison of tonotopic organization with histological data and literature

Figure 9A and G show two relatively superficial brain sections (within 80 μm and 120 μm from dorsal surface, respectively) from one Gad2;Ai96 and one GP4.3 animal stained for GAD67 after two photon imaging. We overlaid the line representing the neurons with the lowest CF and compared its location with the border of LCIC and DCIC traced from the latest Allen Reference Atlas (CCFv3) and from the classical reference atlas by Paxinos and Franklin (2001). The minimum frequency line aligned with the demarcation from the Allen Reference Atlas, while that by Paxinos and Franklin lied in the orthogonal direction.

**Figure 9.**
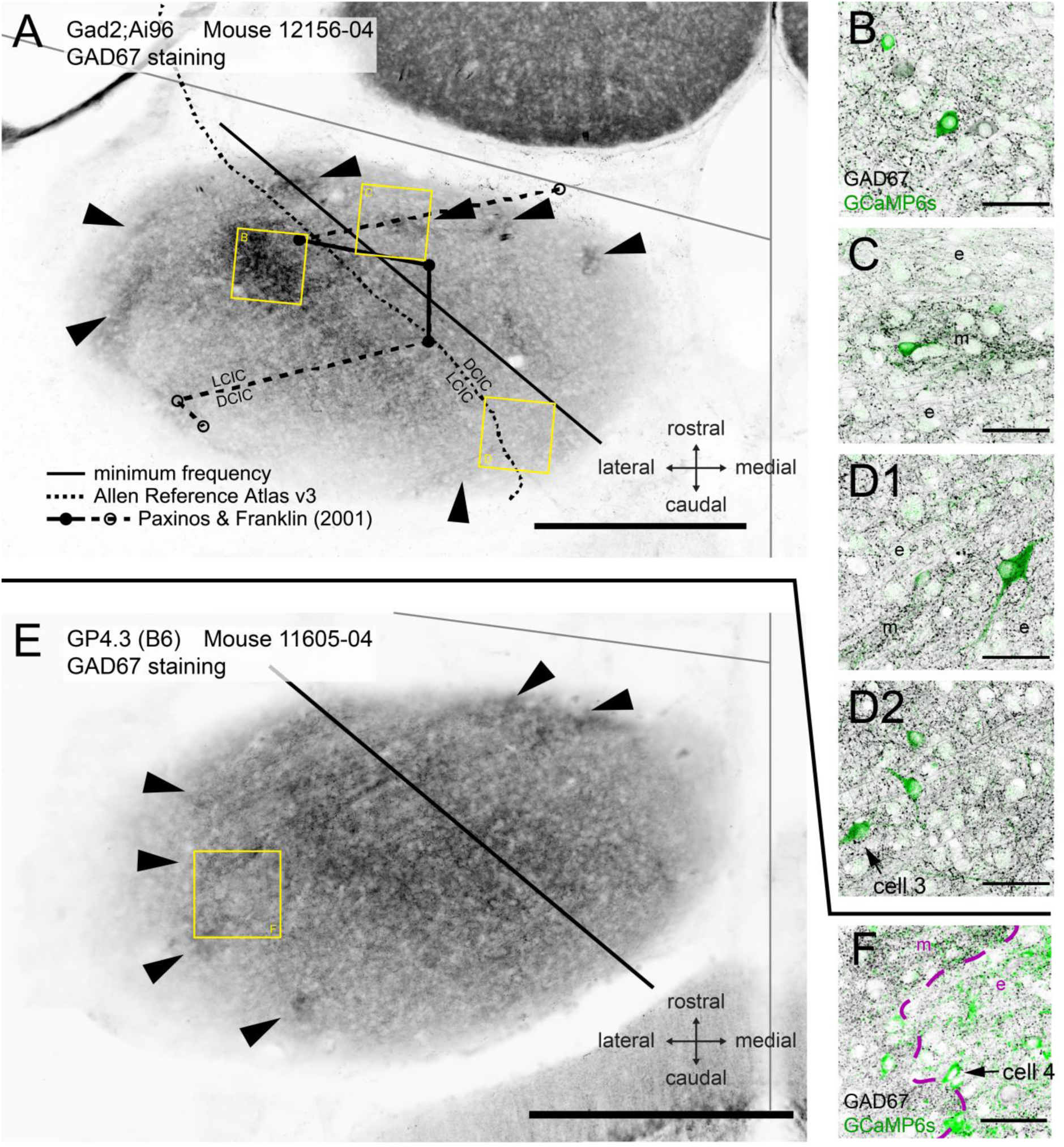
Comparison of tonotopic organization with histological data and literature. (A) Epifluorescence image of the IC in a horizontal brain section stained for GAD67. This brain slice was from a Gad2;Ai96 mouse after two-photon imaging. Black straight line indicates the minimum frequency location derived from fitting two-photon imaging data. Dashed curve represents the demarcation between the dorsal and the lateral (external) cortices traced from version 3 of the Allen Reference Atlas. Circles connected with solid and broken lines mark the demarcation at the dorsal brain surface traced from the atlas by Paxinos and Franklin (2001). The rostral and caudal end of this demarcation are marked by a dashed line because at the indicated positions on the anterior-posterior axis the whole structure was labeled as LCIC or DCIC, respectively. Modules with dense GAD67 staining, considered to be a hallmark for the LCIC, were observed both medially and laterally from each of the three demarcations (arrowheads). We observed a region in the center of the IC whose GAD67 staining density was at least as strong as in the neurochemical modules (square labelled B). (B-D) Single confocal optical sections of GAD67 (black) and GCaMP6s (green) staining in different areas of the brain slice corresponding to the yellow squares in (A). (B) Dense GAD67 area in the central region of the IC (<80 µm from dorsal surface). (C) An area showing a GAD67-dense module (m) in the center, cut transversely and surrounded by extramodular region (e) with sparse GAD67 staining. (D1-2) Optical sections at different focal depths of an area showing another GAD67 module from the same slice (m), cut tangentially. Arrow in D2 indicates a cell showing motion-related responses. (E) Similar to A but from an imaged GP4.3 mouse. (F) Single confocal section for GCaMP6s (green) and GAD67 (black) staining in region marked in E. Arrow marks another cell with motion-related response. Scale bars, A,E: 500 μm; B-D,F: 50 μm.

Neurochemical modules with a high density of GAD67-positive terminals have been reported as a hallmark for the LCIC (Chernock et al. 2004; Lesicko et al. 2016; Dillingham et al. 2017, Video 2). These modules are the predominant target of the somatosensory projections from the cerebral cortex as well as other brainstem areas (Lesicko et al. 2016). We also observed a central strip of dense GABAergic staining, which was not contiguous with the modules (Figure 9 – figure supplements 1-3).

How do motion-sensitive cells associate with the GAD67-dense modules? Figure 9D2 and F show two motion-sensitive neurons retrieved in histology (cells 3 & 4, marked by arrows), both of which resided in close proximity to a GAD67-dense module. We observed that although the somata of both cells resided in the extramodular regions, both possessed dendrites that extended into a GAD67-dense module (Figure 10). Figure 10A shows a dendritic extension from cell 3. For cell 4, a pixel-wise correlation of the *in vivo* somatic fluorescence (Junek et al. 2009) change nicely revealed its dendritic arbor at the same focal plane (Figure 10E). Aligning this to *post hoc* histological staining (Figure 10D), we can demonstrate that cell 4 has dendritic branches extending both inside and outside of GAD67 modules (Figure 10F). We suggest that this may be a functional connection scheme for extramodular neurons to enable them to integrate somatosensory and auditory inputs to the IC (Figure 10G).

**Figure 10.**
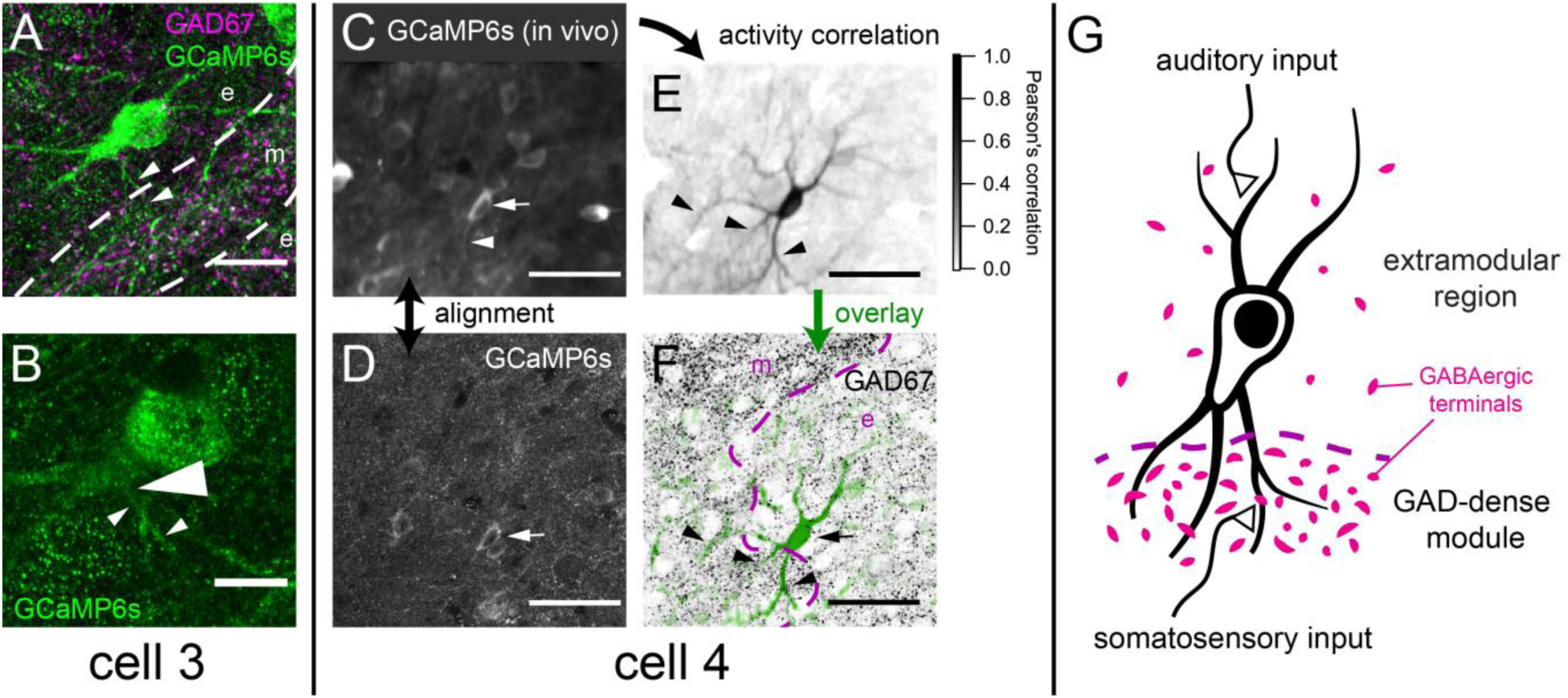
Example motion-sensitive cells with dendritic arbor extended into GAD67-dense modules. (A) Maximum projection of GCaMP staining showing dendritic arbor of cell 3 in Figure 9D2, overlaid with single optical section of GAD67 staining showing a module (m; at same focus as Figure 9D2). While its soma is extramodular (e), at least one branch of its dendrite (arrowheads) appeared to extend into the modular region. (B) A single confocal section of cell 3 showing the root (large arrowhead) of the dendrite labelled in A (small arrowheads). (C) Averaged GCaMP6s fluorescence for the *in vivo* session imaging area in Figure 9F, revealing a dendrite of cell 4 extending into the modular region. (D) Single confocal section for GCaMP6s in fixed brain slices. (E) Dendritic arbor of cell 4 was revealed using pixel-wise correlation to the average somatic fluorescence (Junek et al. 2009), showing extension into the modular region (arrowheads). (F) Background subtracted pixel correlation from E (green) overlaid onto GAD67 staining (black). (G) Schematic representation of hypothesis that integration of auditory and non-auditory inputs by multisensory neurons in the IC can be based on extension of their dendrites into both modular and extramodular regions. Scale bars: A: 20 μm; B: 10 μm; C-F: 50 μm.

## Discussion

We investigated the functional organization of the dorsal IC in awake mice at the single neuron level. We found that the dorsal IC is tonotopically organized, with isofrequency bands running in a rostrolateral-caudomedial orientation with frequency tuning varying along the caudolateral-rostromedial direction. The lowest CF band was found in the middle, dividing the dorsal IC into two reversed tonotopic gradients. In the caudolateral part, but not in the rostromedial part, neurons that were sensitive to whisker or other movements were found, which was in agreement with the view that the caudolateral and rostromedial part corresponded to the LCIC and DCIC, respectively. We observed four different types of firing patterns in response to tones, but other than the tonotopical organization, no obvious topographical organization was observed for the firing patterns. GABAergic neurons were on average somewhat larger, but otherwise there were not clear differences with glutamatergic neurons in spatial organization or firing patterns. Our experiments thus provide a functional definition of the major organization principles of the dorsal IC in adult mice.

### Suitability of GP4.3 and GAD2-cre mice for 2P imaging in awake mice

Because of the prominence of GABAergic neurons and their substantial contribution to ascending projections (Schofield and Beebe 2019), we compared the responses and functional organization of GABAergic and glutamatergic neurons in the dorsal IC using in vivo two-photon calcium imaging. We confirmed that virtually all neurons expressing GCaMP6s in the GP4.3 line were glutamatergic, but not all glutamatergic neurons were labeled. There may have been a preponderance of labeled cells around the modules in the LCIC (Figure 1D), but a systematic screen of markers would be needed to test for a selective enrichment of glutamatergic subtypes in the GP4.3 line. The large majority of GCaMP6s-positive neurons in the IC of mice from the Gad2;Ai96 line expressed GAD67, in agreement with findings in a related mouse line (Gay et al. 2018). In our hands, these two transgenic lines had the advantage that the kinetics of the responses were relatively homogeneous and well approximated by a linear model (Figure 5B), suggesting that the expression levels, which are a major determinant of the Ca^2+^ kinetics (e.g. (Éltes et al. 2019), were similar across cells. Moreover, in contrast to pilot experiments in which we expressed GCaMP6 using AAV, we found little evidence for toxicity. The more sparse expression facilitated the isolation of the responses of individual neurons.

A substantial fraction of cells did not respond to tones. Low probability responses may have been below detection, since the juxtacellular recordings indicated that single APs could not always be detected. Some cells may respond preferentially to more complex sound stimuli, as previously found for neurons in the shell region (Ehret and Moffat 1985; Aitkin et al. 1994). As discussed below, some cells may prefer somatosensory (or motor) stimuli instead of sound stimuli.

### Response types

In both transgenic lines, we observed four different tone-evoked fluorescence response patterns in awake mice: onset, sustained, offset and inhibitory. Our simultaneous juxtacellular recordings showed corresponding patterns. These types were typically also observed in several earlier electrophysiological studies (e.g. Willott et al. 1988a; b; Jain and Shore 2006; Xie et al. 2007).

The observed four kinetic response types were present in both GABAergic and glutamatergic IC neurons. Apart from a possibly lower fraction of GABAergic neurons with pure offset responses (Figure 4B), our results are thus in line with a recent study that showed that GABAergic and glutamatergic IC neurons have similar response properties throughout the IC (Ono et al. 2017).

The prominence of inhibitory responses in the IC (42-52% of sound-responsive neurons) was a striking finding in our recordings. An important factor contributing to their prominence is probably that our experiments were done in awake animals, as a recent single unit recording study showed the presence of inhibitory responses as well as higher spontaneous spike rates in IC neurons of awake mice compared to mice under urethane anesthesia (Duque and Malmierca 2015), and synaptic inhibition has also been shown to be prominently present in whole-cell recordings in awake bats (Xie et al. 2007).

Offset responses can be found throughout the auditory system including the IC, and they may have an important role in perceptual grouping or in duration discrimination (Kopp-Scheinpflug et al. 2018). However, studies in the IC of anesthetized mice, rats and chinchillas showed that offset responses are generally restricted to cells showing band-pass duration tuning, which only responded to much shorter tones than the 1 second employed here (Chen 1998; Brand et al. 2000; Pérez-González et al. 2006). Another striking finding was therefore that we saw clear evidence for offset responses in 3-11% of sound-responsive cells. A whole-cell study in the mouse IC showed that in most cases offset responses are inherited from upstream, but that they may also be generated *de novo* as a rebound from inhibition (Kasai et al. 2012). Similar to the inhibition class, we therefore suggest that the prominence of offset responses may be related to the increased impact of synaptic inhibition in the IC of awake animals. A study using communication calls suggests that offset responses may be even more prominent following more complex sound stimuli, such as communication calls (Akimov et al. 2017), possibly by a summation of inhibition at different frequency bands (Sanchez et al. 2008).

### Tonotopical organization of the dorsal IC

We observed a tonotopic gradient that ran along a gradient from caudolateral to rostromedial. The gradient reversed at the site where the CFs were the lowest. This reversal is in line with the banding patterns observed with epifluorescent calcium imaging in unanesthetized mice before hearing onset; neurons in these bands fire together, and this activity depended on the cochlea, suggesting that they represented groups of cells with similar CF (Babola et al. 2018). Interestingly, our data are compatible with two earlier calcium imaging studies, even though in the first, which focused on medial regions of the dorsal IC, the gradients were reported to run from lateral (low-frequency) to medial (high-frequency) (Ito et al. 2014), whereas in the second they were reported in a roughly (medio)rostral (low-frequency) to (latero)caudal (high-frequency) direction (Barnstedt et al. 2015). Following pooling of the data from several experiments, we observed a gradient reversal, which, interestingly, was also observed in one animal in the latter study (Barnstedt et al. 2015).

We found considerable variability in CFs within a band. Imprecisions in the alignment of imaging areas and variability between animals may have contributed to the variability, but substantial variability was also observed within a single animal, in line with previous results (Ito et al. 2014; Barnstedt et al. 2015). Two-photon calcium imaging studies in the mouse auditory cortex have met with variable amount of microheterogeneity, apparently depending on layer, calcium dye, or the use of anesthetics (Bandyopadhyay et al. 2010; Rothschild et al. 2010; Winkowski and Kanold 2013; Issa et al. 2014; Kato et al. 2017; Tischbirek et al. 2019).

The four response types showed a similar tonotopic organization, but considerable microheterogeneity. This microheterogeneity was larger than in an earlier study in which bulk loading of an organic Ca indicator was used (Ito et al. 2014), which may have made it more difficult to isolate responses from individual cells. In the auditory cortex, on and off cells showed a similar but not identical tonotopic organization. Our results are in line with patch-clamp results showing considerable heterogeneity between adjacent cells within the dorsal IC, which extended to heterogeneity in their inputs (Geis et al. 2011). The microheterogeneity was similar for GABAergic and glutamatergic neurons. Our results thus differ from auditory cortex, where response patterns are different and local microheterogeneity is much smaller for (parvalbumin-positive) GABAergic neurons than for glutamatergic neurons (Maor et al. 2016; Liang et al. 2018; Liu et al. 2019). We conclude that despite the shared input and firing behavior during development (Babola et al. 2018), functional heterogeneity dominates within bands in the dorsal IC of adult mice. Whereas some of the mechanisms that underlie the microheterogeneity within the auditory cortex are being elucidated (Kato et al. 2017; Tao et al. 2017; Vasquez-Lopez et al. 2017), they remain to be explored for the dorsal IC.

### Presence of somatosensory inputs in LCIC

We monitored spontaneous whisker and general body movements, and used strict criteria to exclude movement artefacts. As we restricted the analysis to neurons that were not excited by tones, we consider it unlikely that the observed somatosensory responses were instead caused by self-generated sounds. Despite these severe restrictions, we did find that a few percent of the cells were excited during whisker or body movements, suggesting a substantial role for somatosensory (or motor) inputs, especially considering that in previous research only a minority of cells have been shown to respond to unimodal tactile stimuli, and the dominant effect of somatosensory inputs appears to be inhibitory (Aitkin et al. 1978; Aitkin et al. 1981; Zhou and Shore 2006). We did not further discriminate between different movements as they tended to be heavily correlated. Moreover, it is known that neurons in the LCIC have broad tuning for somatosensory input (Aitkin et al. 1981). Movement-sensitive neurons were not found in the high-frequency DCIC regions, but only in the putative CNIC region and the LCIC.

A defining feature of the LCIC is the presence of neurochemical modules, which have been shown in many species, including rats (Chernock et al. 2004; Choy Buentello et al. 2015), but also in both adult (Choy Buentello et al. 2015; Lesicko et al. 2016; Patel et al. 2017) and developing mice (Dillingham et al. 2017). While it is still unclear how neurons in these modules and extramodular zones connect with each other, there is evidence that somatosensory inputs project predominantly to the modules, while auditory input projects to the extramodular region (Lesicko et al. 2016). The main sources for somatosensory input to the LCIC are the spinal trigeminal nucleus, the dorsal column nuclei and the somatosensory cortex (reviewed in Gruters and Groh 2012). These inputs target the layer 2 modules, which constitute only a small fraction of the LCIC volume. Moreover, neurons in the modules do not seem to target extramodular IC neurons (Lesicko and Llano 2019). This raises the question how so many cells in the LCIC can be sensitive to somatosensory inputs. We obtained anecdotal evidence that neurons at the border of the modules can extend dendrites into the modules (Figure 10A,F), which would allow them to sample somatosensory inputs. Indeed, neurons in LCIC can have extensive dendritic trees (Smith 1992), and Golgi stainings show that cells in layer 3 of LCIC can send dendrites up to layer 1 (Meininger et al. 1986; Malmierca et al. 2011). The CNIC has few somatosensory inputs, but at least half of the cells in CNIC are affected by stimulation of the dorsal column nuclei (Gruters and Groh 2012). The CNIC may get somatosensory inputs indirectly via the dorsal cochlear nucleus, which is innervated by the spinal trigeminal nucleus and the dorsal column nuclei, or in the form of intracollicular input from the LCIC.

### Functional parcellation of the dorsal IC

There is little agreement on the precise borders of the different nuclei of the IC, and its parcellation has been different based on whether cell morphology, inputs, or physiological properties were chosen as the main classifier (Oliver 2005). Here we used physiological properties as the main criterion to look at the borders between the DCIC, LCIC and CNIC in the dorsal IC. Our findings suggest that the laterocaudal area with high frequencies is the dorsal edge of the LCIC, whereas the mediorostral part of the dorsal IC belongs to the DCIC. The tonotopic reversal would thus provide a functional demarcation between the dorsal and lateral cortices of the IC. Earlier tracing studies of intracollicular projections in the guinea pig (Malmierca et al. 1995) and the rat (Saldaňa and Merchán 1992), as well as ascending projections from the cochlear nucleus in the rat (Malmierca et al. 2002), have shown interesting V-shaped arrangements in coronal sections, where the medial side corresponds to isofrequency laminae in CNIC and DCIC, while the lateral side corresponds to those in the LCIC. These V-shaped axon terminal plexuses, if extended to the dorsal surface of the IC, provide an anatomical explanation for the observed tonotopic reversal. This is also in agreement with the latest anatomical framework published by the Allen Brain Institute (Figure 9A; Allen Mouse Common Coordinate Framework, 2015), previous imaging studies (Ito et al. 2014; Barnstedt et al. 2015; Babola et al. 2018), the known somatosensory inputs to the LCIC, and both the descending and ascending inputs to the IC (Aitkin et al. 1981; Künzle 1998; Jain and Shore 2006; Zhou and Shore 2006; Lesicko et al. 2016; Patel et al. 2017). Surprisingly, we observed GAD67-dense modules in both the medial and the lateral aspect of the IC (arrowheads Figure 9A,G), i.e. within both LCIC and DCIC in either atlas. The medial modules could not be imaged within this study, so their association with somatosensory inputs is currently unclear.

The central strip with neurons tuned to low frequencies in between the LCIC and the DCIC may be the most dorsal extension of the dorsolateral low frequency area of the CNIC (Stiebler and Ehret 1985; Barnstedt et al. 2015). This strip contains the large density of GABAergic inputs (Figure 9A,B) that is characteristic of the CNIC (Choy Buentello et al. 2015). Lemniscal inputs are known to extend quite dorsally, almost up to the surface, and neurons with short latency sound responses have been found in the dorsal IC, although their exact location was not studied (Geis and Borst 2013). The low tuning in the central strip extended close to the surface, but a more systematic study of their properties would be needed to be able to assign the most superficial layers to either the DCIC or another region. From our data it thus appears that the three main areas of the IC, LCIC, CNIC, DCIC, are readily accessible at the dorsal IC. Their accessibility for imaging studies will thus help to further delineate their functions and the role of their inputs in the future.

## Materials and Methods

### Animals

Detailed two-photon imaging experiments were conducted on seven GP4.3 transgenic animals (Dana et al. 2014) (three in C57BL/6J background and four in B6CBAF1/J background) and six F1 progeny between Gad2-IRES-cre (Taniguchi et al. 2011), and Ai96 reporter line (B6;129S6-*Gt(ROSA)26Sor^tm96(CAG-GCaMP6s)Hze^*/J) (Madisen et al. 2015). We will refer to the Gad2-IRES-Cre × Ai96 cross as Gad2;Ai96. Postnatal age at recordings ranged between 11 – 35 weeks (median: 19; Q1: 15; Q3: 23). Ground-truth juxtacellular recordings were performed on four GP4.3 and three Gad2;Ai96 animals. Immunohistochemistry for cell counting was performed on three GP4.3 and three Gad2;Ai96 animals.

GP4.3 animals were originally obtained from the Jackson Laboratory (C57BL/6J-Tg(Thy1-GCaMP6s)GP4.3Dkim/J; JAX stock #024275); they were maintained in a heterozygous state by backcrossing to C57BL/6J from Charles Rivers (JAX™ C57BL/6J). To create GP4.3 animals with B6CBAF1/J background, heterozygous GP4.3 in C57BL/6J background were crossed with CBA/JRj mice from Janvier. Gad2-IRES-Cre (originally STOCK *Gad2^tm2(cre)Zjh^*/J, the Jackson Laboratory) was maintained in homozygous state after >10 generations of backcrossing to C57BL/6J, and re-backcrossed to C57BL/6J every 4-5 generations. Ai96 mice were obtained from Jackson Laboratory already with 3 generations of backcross to C57BL/6J (N3), and subsequently maintained by backcrossing to C57BL/6J for 5-7 generations. All experiments complied with the ethical guidelines for laboratory animals within our institute and with European guidelines, and were approved by the animal ethical committee of the Erasmus MC.

### Surgery

Animals were anaesthetised through respiratory intake of isoflurane and maintained at surgical level of anaesthesia, assessed through the hind limb withdrawal reflex. A heating pad with rectal feedback probe (40-90-8C; FHC, Bowdoinham, ME, USA) was used to maintain body core temperature at 36-37 °C. Eye ointment (Duratears; Alcon Nederland, Gorinchem, The Netherlands) was used to keep the eyes moist during surgery. A bolus of buprenorphine (0.05 mg/kg; Temgesic, Merck Sharp & Dohme, Inc., Kenilworth, NJ, USA) was injected subcutaneously at the beginning of surgery. The skin overlying the IC was incised. Lidocaine (Xylocaine 10%; AstraZeneca, Zoetermeer, The Netherlands) was applied before removing the periosteum and cleaning the skull. After etching the bone surface with phosphoric acid gel (Etch Rite™; Pulpdent Corporation, Watertown, MA, USA), a titanium head plate was glued to the cleaned bone above the left IC using dental adhesive (OptiBond FL; Kerr Italia S.r.l., Scafati, SA, Italy) and further secured with dental composite (Charisma®; Heraeus Kulzer GmbH, Hanau, Germany).

Through an opening in the head plate, a craniotomy of 3 mm diameter centered at one of the ICs was made by thinning and removing the skull bone. A cranial window, made by gluing a 3 mm cover slip (CS-3R-0; Warner Instrument Inc, Hamden, CT, USA) on a custom built, 500 µm thick steel ring with UV-cured optical adhesive (NOA68; Norland Products), was installed over the exposed brain surface and secured with superglue. Each animal was allowed to recover for at least two days before the first measurements. For in vivo electrophysiology, the cranial window construct was gently removed, and the dura mater covering the IC and part of the cerebellum was carefully punctured and removed with a pair of fine forceps.

After all recordings had been done, animals received an intraperitoneal injection of pentobarbital (300 mg/kg) and were perfused transcardially, first with physiological saline (Baxter Healthcare, Zurich, Switzerland), followed by 4% paraformaldehyde (PFA) in 0.1 M phosphate buffer (PB; 4% PFA in PB, pH 7.4; Merck).

### Two-photon imaging

For two-photon imaging of the IC, a 20X water-immersion objective (LUMPlanFI/IR, 20X, NA: 0.95; Olympus Corporation, Tokyo, Japan) on a custom-built two-photon microscope was used, except for simultaneous juxtacellular recordings, for which a long working distance 40x objective (LUMPlanFl/IR, 40X, NA: 0.80, Olympus Corporation, Tokyo, Japan) was used. Excitation light was provided by a MaiTai Ti:Sapphire laser (Spectra Physics Lasers, Mountain View, CA, USA) tuned to a wavelength of 920 nm. A layer of Ringer solution was put between the objective and the cranial window. GCaMP6s fluorescence was captured by a photomultiplier tube (H6780-20, Hamamatsu, Japan) after a barrier filter at 720 nm (FF01-720/SP-25; Semrock), a secondary dichroic at 558 nm and a green bandpass filter centered at 510 nm (bandwidth: 84 nm; FF01-510/84-25; Semrock); AlexaFluor 594 fluorescence was captured at the second channel with a red bandpass filter centered at 630 nm (bandwidth: 60 nm; D630/60; Chroma). Images (256×128 pixels) were collected at 9 Hz (2 μs/pixel; 1-2 μm/pixel). To minimize acoustic noise from scanning, a sinusoidal waveform was used for the X Galvo-scanner. Data were acquired in the middle 80% of the sinusoidal waveform to minimize nonlinearity. Multiple regions were imaged sequentially in multiple sessions in awake, head-fixed animals. For depth estimation, a Z-stack was acquired over a 512×512 µm area at 1×1×1 µm voxel size. Depth of each imaged area was estimated by measuring the Z-distance from the pia surface. Neurons in this dataset lied between 15 – 155 μm from the pia surface. The relative position of each region was tracked through the micromanipulator (MP-285, Sutter Instrument) that controlled the microscope objective. The positions of imaged areas were further aligned across animals to a common coordinate using the midline, lateral extreme and anterior-posterior extremes of the IC as anatomical landmarks in bright field images (SZ 61,Olympus) of the cranial window.

Frame timing of the scanner, timing of the sound stimuli, and animal movements were digitized using a Digidata 1440A (Molecular Devices, Sunnyvale, CA, USA) with Clampex v. 10.3 (Molecular Devices, Sunnyvale, CA, USA).

### Juxtacellular Recording

*In vivo* juxtacellular recordings were made under two-photon guidance (Kitamura et al., 2008). Glass pipettes were pulled from 1.5 mm, thick-walled borosilicate capillaries (Hilgenberg, Malsfeld, Germany) to 1–2 µm tip diameter (P-97; Sutter Instrument, Novato, CA) and filled with internal solution containing (in mM): potassium gluconate 138, KCl 8, Na_2_-phosphocreatine 10, Mg-ATP 4, Na_2_-GTP 0.3, EGTA 0.5, HEPES 10, (pH 7.2 with KOH; Merck). The internal solution also contained 40 µM Alexa Fluor 594 hydrazide. Ringer solution containing 1-2% agarose and (in mM): NaCl 135, KCl 5.4, MgCl_2_ 1, CaCl_2_ 1.8, HEPES 5, was applied on the brain surface to reduce movement artefacts. A positive pressure of about 300 mbar was maintained before penetration of the pia surface, and reduced to about 30 mbar upon pia entry; pressure was removed upon cell approach. Electrode resistances were constantly monitored and recording started when resistance increased to >25 MΩ. Juxtacellular potentials were acquired with a MultiClamp 700A amplifier (Molecular Devices, Sunnyvale, CA, USA) in current-clamp mode. Signals were low-pass filtered at 10 kHz (four-pole Bessel filter) and digitized at 25 kHz (Digidata 1322A). Data were recorded with pCLAMP 9.2 (Molecular Devices). In some cases a large current injection (1-6 nA) was used to elicit (positive current) or suppress (negative current) spikes of the cell being recorded (nanostimulation; e.g. Houweling et al. 2010), usually for cells that did not show obvious change in spike rate upon pure-tone stimulations. These data were also used for fitting the fluorescence ground-truth model.

### Sound stimulation

Tone stimuli were generated in MATLAB v7.6.0 (The MathWorks, Natick, MA, USA) and played back via a TDT System3 setup (RX6 processor, PA5 attenuator, ED1 electrostatic speaker driver and two EC1 electrostatic speakers; Tucker Davis Technologies, Alachua, FL, USA). Sound stimuli were presented bilaterally in open field. Sound intensities were calibrated using a condenser microphone (ACO pacific Type 7017; ACO Pacific, Inc., Belmont, CA, USA) connected to a calibrated pre-amplifier and placed at the position of the pinnae.

For measurement of frequency response areas (FRA), 1 s tones (including 2.5 ms cosine-squared rise/decay times) with frequencies between 1 and 64 kHz with three steps per octave were presented at intensities between 30 and 80 dB sound pressure level (dB SPL) in steps of 10 dB. The set of stimuli was presented 6-10 times per experiment each in a pseudorandom order at an inter-stimulus interval of 1.5 s. For juxtacellular recordings, tones were only presented at one or two intensities (70 dB SPL; or 50 & 70 dB SPL) due to the limited recording time.

### Behavioural measurements

Visual recordings were made using an RS Miniature CCD Camera (RS Components, Corby, UK) at 3 Hz. The camera was aimed at the animal’s head from a roughly right lateral perspective, in order to clearly record the animal’s right eye and whiskers. Facial movement was detected by calculating the root-mean-square intensity change between successive frames in a rectangular region at the whisker pad. General movement of the animal was registered using a piezo-electric motion sensor under the front paws of the mouse, digitized without further amplification using an AD channel of the Digidata 1440A.

### Immunohistochemistry and cell counting

Gelatin-embedded, 40 μm sections of PFA-fixed mouse brains were stained with the following antibodies: mouse anti-Gad67 (cat. no.: MAB5406; Millipore; 1:1000; RRID: AB_2278725); chicken anti-GFP (cat. no.: GFP-1020; Aves; 1:1000; RRID: AB_10000240); rabbit anti-NeuN (cat. no.: ABN78; Millipore; 1:1000; RRID: AB_10807945); mouse anti-parvalbumin (cat. no.: 235; Swant; 1:7000; RRID: AB_10000343); rabbit anti-Calretinin (cat. no.: 7699/4; Swant; 1:5000; RRID: AB_2313763); and AlexaFluor-, Cy3- or Cy5-conjugated secondary antibodies (Invitrogen or Jackson ImmunoResearch). To ensure a more homogeneous GAD67 fluorescence for the *post-hoc* immunostaining of imaged brains, the brain slices were incubated twice in the primary antibody solution, each for 1 week at 4 °C. The secondary antibody was also applied twice, but overnight at room temperature. Cell counting was performed manually in FIJI using the Cell Counter plug-in on confocal z-stacks.

### General Analysis

Data analysis was performed with Igor Pro (WaveMetrics, Inc., Lake Oswego, OR, USA) using custom written procedures. Two-photon images and behavior video were aligned to ClampEx data using stimulus timing. Whisking behavior was assessed by calculating the root-mean-square of the frame to frame intensity difference in an area at the whisker pad.

Movement artefacts in two-photon images were corrected based on the built-in ImageRegistration operation in Igor Pro, which is based on a published algorithm (Thevenaz et al. 1998). Neuronal cell bodies were identified visually based on the average image of the motion-corrected image series, and a higher sampling Z-stack of the area (0.5×0.5 µm pixels; 1 µm z steps) taken directly after each experiment. Regions-of-interests (ROIs) were drawn around cell bodies, and an average fluorescence was extracted for each ROI after subtracting the average fluorescence fluctuation of the surrounding background for further analysis. The background fluorescence was defined as the average fluorescence in a 2 µm wide contour surrounding the ROI, excluding any pixel directly belonging to another ROI.

### Analysis of Frequency Response Areas

The frequency response area (FRA) of each ROI was analyzed largely as described previously (Geis *et al*., 2011). The fluorescence trace within 1 s of each stimulus was taken as the stimulus related waveform. To extract the stimulus-related response, we extracted the *signal autocorrelation* (also named “consistency index” by Barnstedt *et al*., 2015) by calculating the average Pearson correlation coefficient among fluorescence waveforms to the same stimuli.

Kinetics of sound-evoked responses were fitted by single exponential functions.

### Orientation of CF Gradient

To find the most prominent direction of CF gradient, we parametrized the location of each neuron as a projected distance *r* from the anatomical origin along a line with an angle θ with the medial-lateral orientation:

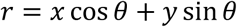

where x & y are coordinates (in micrometers) of cells along the medial-lateral and anterior-posterior axes, respectively. We then model CF as a polynomial function of *r*:

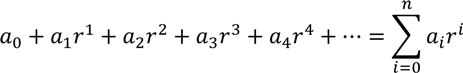

This whole function was then fitted to the logarithm of the CF values of the dataset using the Levenberg-Marquardt least-squares method, implemented in Igor Pro, i.e.:

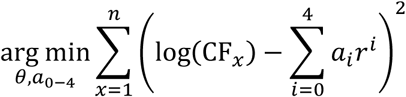

A 4^th^ order polynomial was used in Figure 6 because increasing the number of exponents did not increase the explained variance further. The best fit parameters were θ = 0.875 rad (≈50°), *a*_0_= 6.95, *a*_1_=−0.0135 dec μm^−1^, *a*_2_= 1.95 × 10^−5^ dec μm^−2^, *a*_3_= −1.11 × 10^−8^ dec μm^−3^, *a*_4_= 2.17 × 10^−12^ dec μm^−4^. Cross-validation (70% fitting/30% test) indicated a slight overfitting. We also considered 2D polynomial fits in the form ∑ aij xi yj where 0 ≤ *i* + *j* ≤ *n*, and found that a 3^rd^ order (*n* = 3) fit would be an optimal compromise between bias (underfitting) and variance (overfitting). However, we opted for the single dimension polynomial to avoid potential overinterpretation of the complex contour created by the 2D polynomial.

### Ground-truth and modelling of GCaMP6s fluorescence

Traces from juxtacellular recording were first subjected to a digital DC remove filter which subtracts at each point in time the average potential within ±1 ms to remove DC drift or offset introduced by nanostimulation (Houweling et al. 2010). Traces were blanked around the start and end of each current injection (2 ms before and 3 ms after) to remove stimulus artefacts. Spikes were then detected by a simple thresholding procedure, with spike times defined as the peak time of the spikes, which presumably corresponds to the maximum rate of rise of the action potential.

For spike vs fluorescence model, spike times were converted to number of spikes over time (*n(t)*) at the same sampling rate of the imaging (114.4 ms bins). It was then convolved with a simple ramp-decay kernel (*g(t)*) according to the equations:

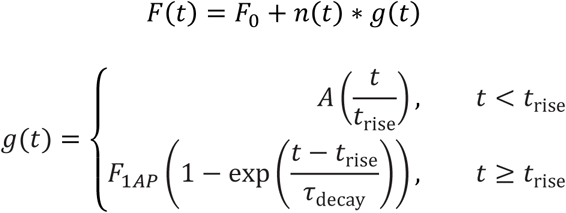

where F_1AP_ is the amplitude of the fluorescence change per action potential, t_rise_ is the rise time of the fluorescence and τ_decay_ is the decay time constant for the fluorescence. An offset (F0) is added, which represents the minimal fluorescence at zero spike rate.

This 4-parameter model was fitted to the fluorescence trace using a genetic fitting routine in Igor Pro (Gencurvefit XOP, kindly provided by Andrew Nelson, Australian Nuclear Science and Technology Organisation).

### Statistics

Statistical significance of signal correlation was done by a bootstrap method with Holm-Bonferroni correction for multiple comparisons: for each ROI, a distribution of average Pearson correlation coefficient was constructed by drawing 30,000 random samples of 10 fluorescence trace segments within an experimental session. The p-value of each stimulus (with *n* repetitions) was calculated as the fraction of *n*-member samples having a greater (for FACA) average Pearson’s correlation than that of the stimulus. The ranked p-values were then tested for significance with α = 0.05 and Holm-Bonferroni correction for the number of different stimuli presented (19 frequencies × 6 intensities = 114 for FRA measurements).

## Acknowledgements

We thank Kees Donkersloot and Elize Haasdijk for their excellent technical support, H.-Rüdiger A.-P. Geis for help in setting up two-photon imaging experiments and Alba Membrilla Esteban for performing part of the immunohistochemistry analysis, Zhenyu Gao for sharing the design of the chronic cranial window, and Travis Babola and Dwight Bergles for helpful discussions on tonotopy. We thank Douglas Kim and the GENIE Project at the Janelia Research Campus for making the GP4.3 mouse line available.

This work was supported by a NeuroBasic PharmaPhenomics grant (AgentschapNL of the Ministry of Health, Welfare and Sports of the Netherlands to JGGB). ABW was supported by an Individual Fellowship through the Marie Skłodowska-Curie actions (European Commissions).

---

**Figure 1 – figure supplement 1.**
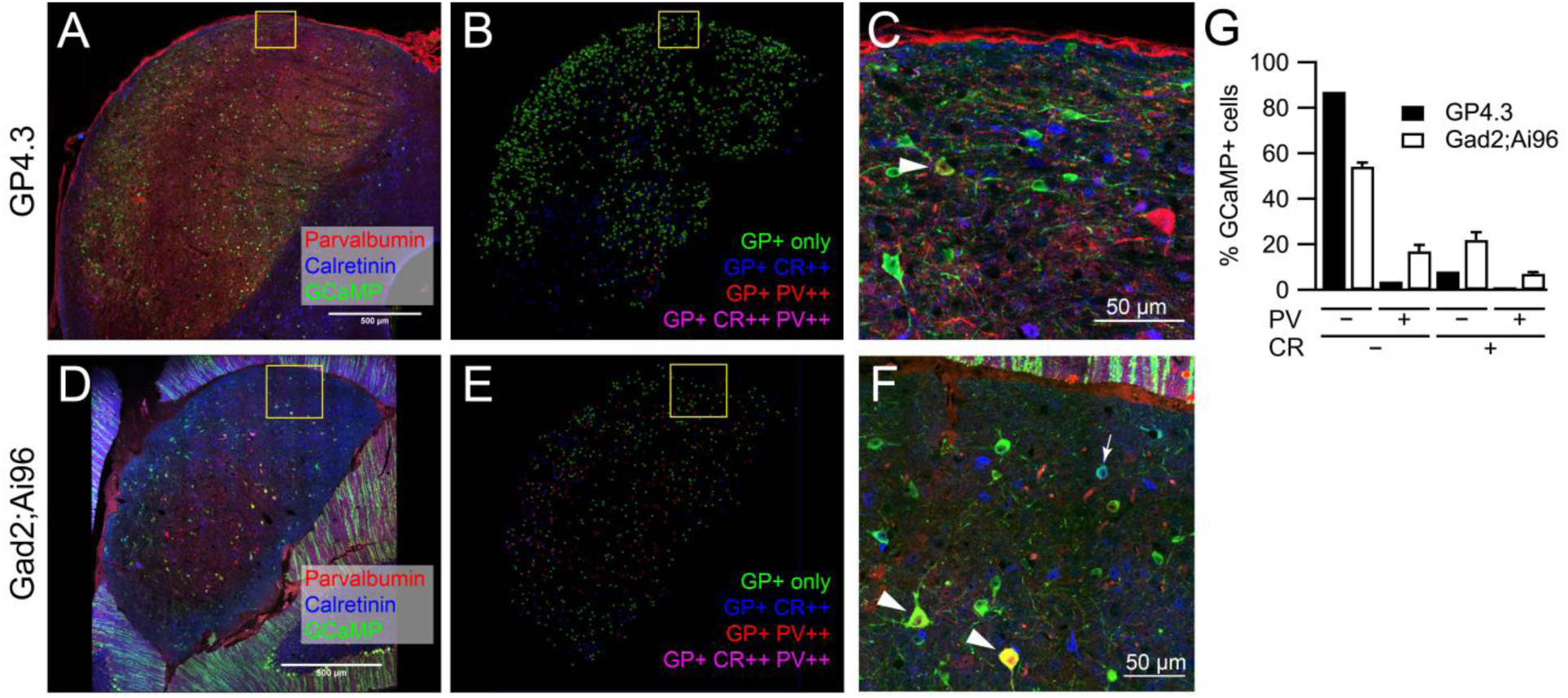
Expression of parvalbumin and calretinin in GCaMP+ neurons in the two transgenic mouse lines. (A) Single optical section of an IC brain slice from a GP4.3 mouse immunolabeled for GCaMP6s (green), parvalbumin (PV; red) and calretinin (CR; blue). (B) Distribution of GCaMP+ cells in the 40 µm brain slice in (A), color-coded by their immunoreactivity to PV and CR antibodies. (C) Enlarged image from A (square), showing different combinations of immunoreactivity. Arrowhead: a GCaMP+CR-PV+ cell. (D,E) As A and B, but from a Gad2;Ai96 mouse. (F) Enlarged region from D (square). Arrowheads: GCaMP+CR-PV+ cells; Arrow: a GCaMP+CR+PV-cell. (error bars: s.e.m.; GP4.3: n = 2 slices from 2 animals; Gad2;Ai96: n = 3 slices from 2 animals).

**Figure 3 – figure supplement 1.**
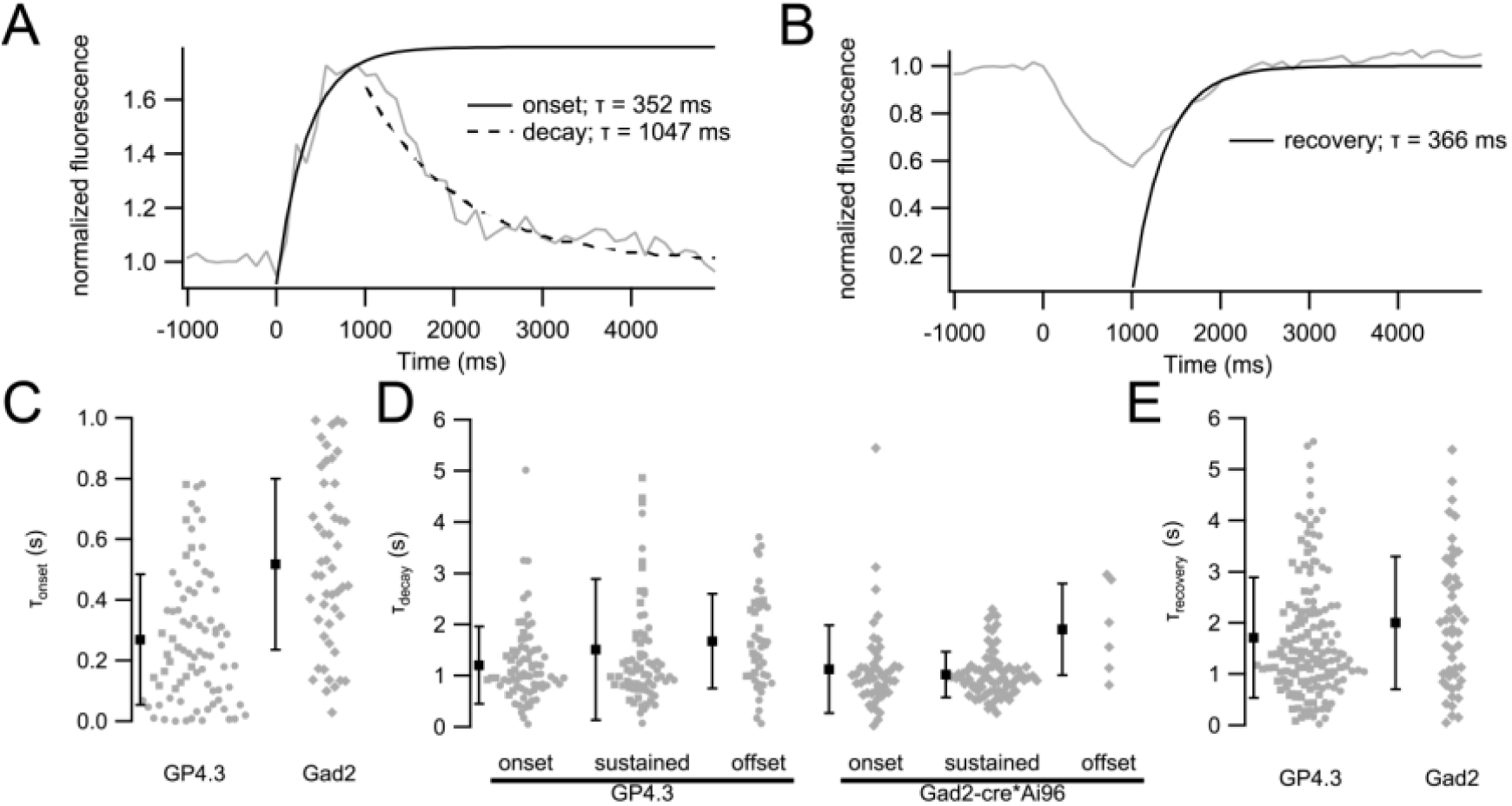
Kinetics of fluorescence responses. (A) Example of the average fluorescence response of a cell with an onset-type response to sound (grey trace). Single exponential fits were performed on the onset (0-1000 ms; solid black curve) and decay (1.5-5 s; dashed black curve) periods. Due to the short inter-stimulus interval (1.5 s), responses where another effective stimulus followed immediately were excluded for the averaging. (B) For inhibitory responses, the recovery from inhibition was used for the fitting (1.5-5 s; solid black curve). (C) Onset time constant for cells showing onset response (τ_onset_) (GP4.3 mean τ_onset_: 269 ± 215 ms, n = 81 cells; Gad2;Ai96 mean τ_onset_ = 518 ± 282 ms, n = 49 cells) (D) Decay time constant (τ_decay_) for cells showing onset, sustained or offset responses. Mean τ_decay_ for onset cells (GP4.3: 1200 ± 756 ms; Gad2;Ai96: 1120 ± 860 ms), sustained cells (GP4.3: 1510 ± 1380 ms; Gad2;Ai96: 1020 ± 440 ms) and offset cells (GP4.3: 1670 ± 920 ms; Gad2;Ai96: 1900 ± 890 ms). (E) Recovery time constant (τ_recovery_) for inhibited cells (GP4.3: mean τ_recovery_: 1710 ± 1180 ms, n = 153 cells; Gad2;Ai96 mean τ_recovery_ = 2000 ± 1300 ms, n = 50 cells). Error bars in C-E represent s.d.

**Figure 3 – figure supplement 2.**
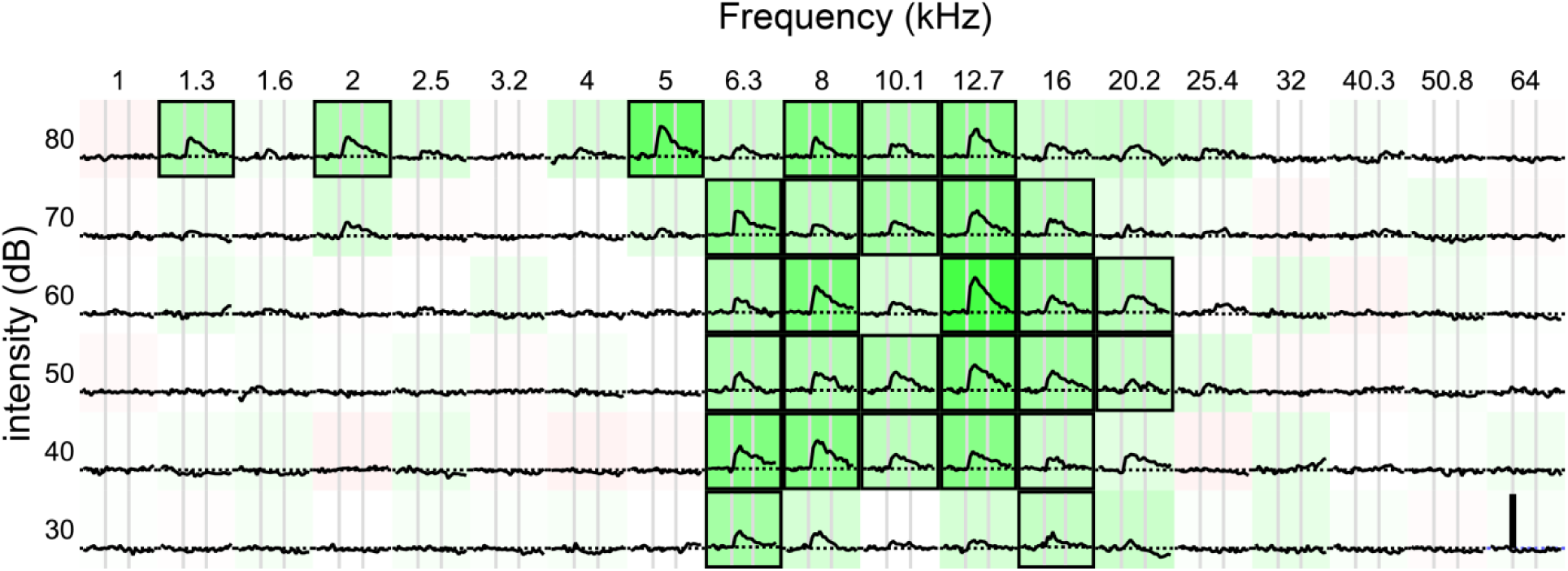
Another example FRA from GP4.3 mouse, showing onset response and broad tuning.

**Figure 8 – figure supplement 1.**
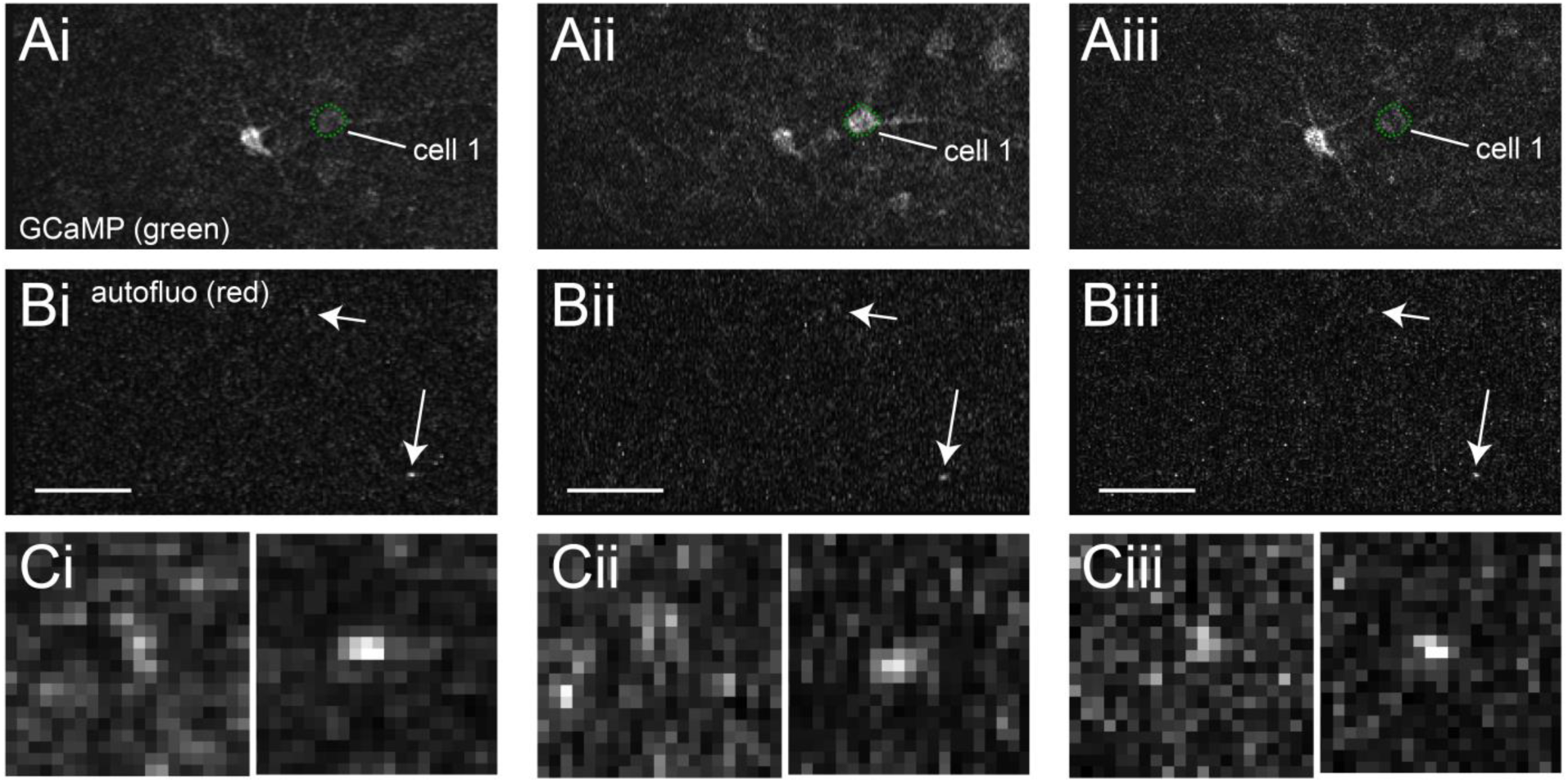
Evidence that the motion-related responses were not due to motion artefacts in imaging. Vertical or horizontal blurring was due to subpixel image registration. (A) Single image frame for GCaMP fluorescence from experiment in Figure 8, i-iii corresponds to frames marked in Figure 8A (B) Autofluorescence in red channel for the same frames in A. Arrows point to two autofluorescence spots, both of which remained in focus during this period. (C) Enlarged image of the two autofluorescence spots for the three frames.

**Figure 9 – figures supplement 1.**
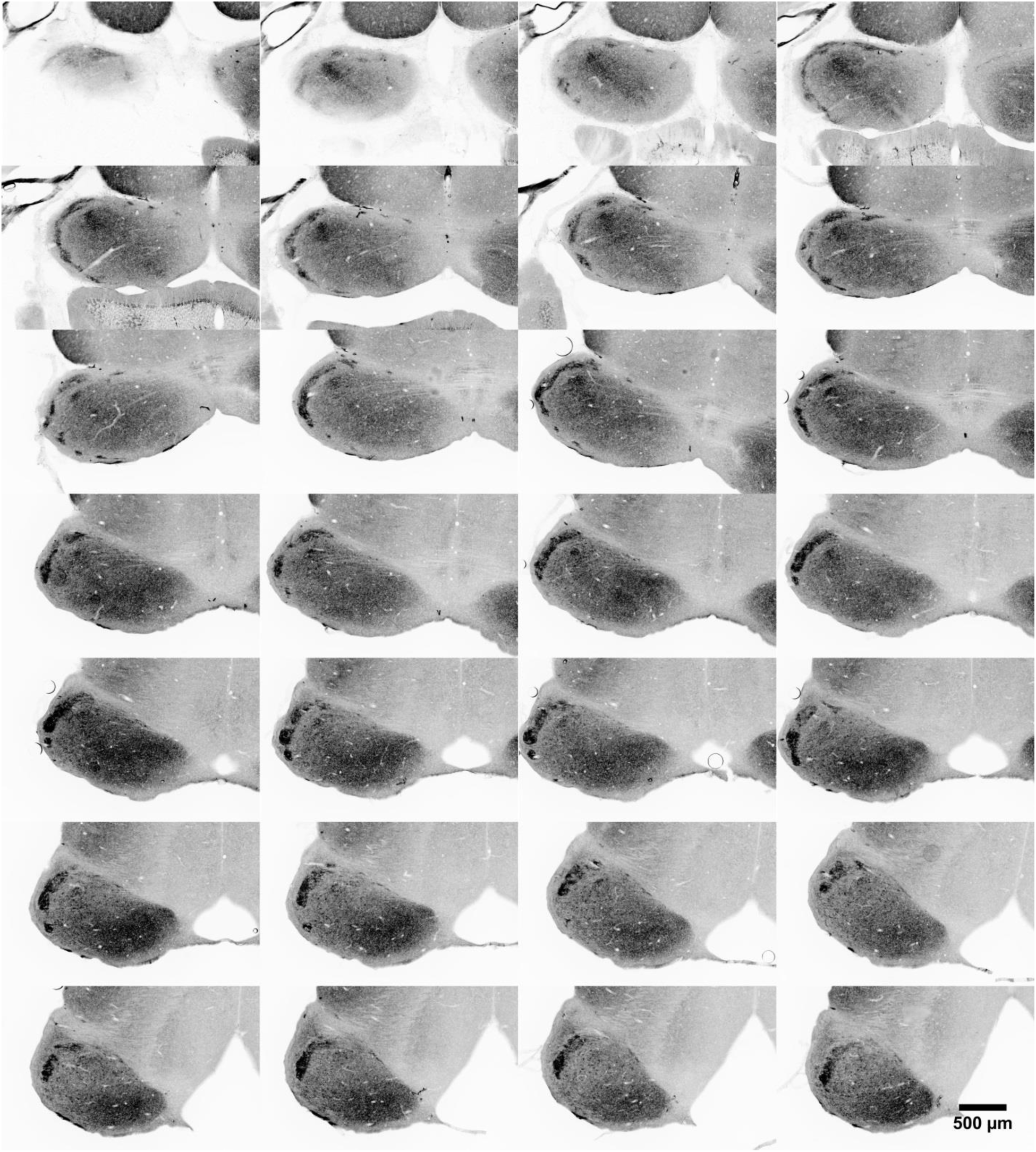
GAD67 staining in the inferior colliculus of the series of consecutive 40 μm horizontal brain slices from the same animal as in Figure 9A, displayed from dorsal (top-left) to ventral (right-bottom).

**Figure 9 – figures supplement 2.**
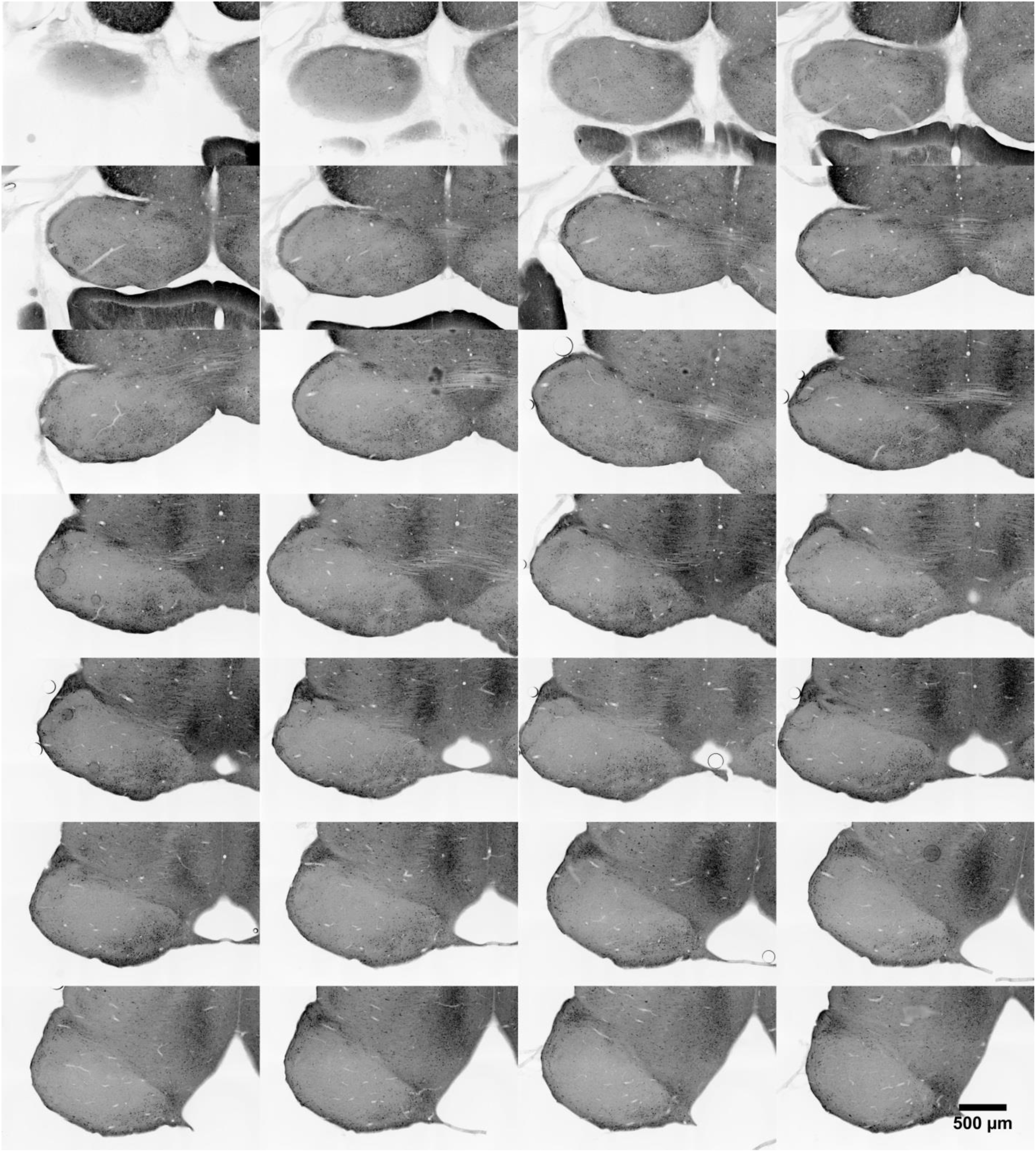
Calretinin staining in the inferior colliculus of the series of consecutive 40 μm horizontal brain slices from the same animal in Figure 9A, displayed from dorsal to ventral (left to right, top to bottom). Same sections as supplement 1.

**Figure 9 – figure supplement 3.**
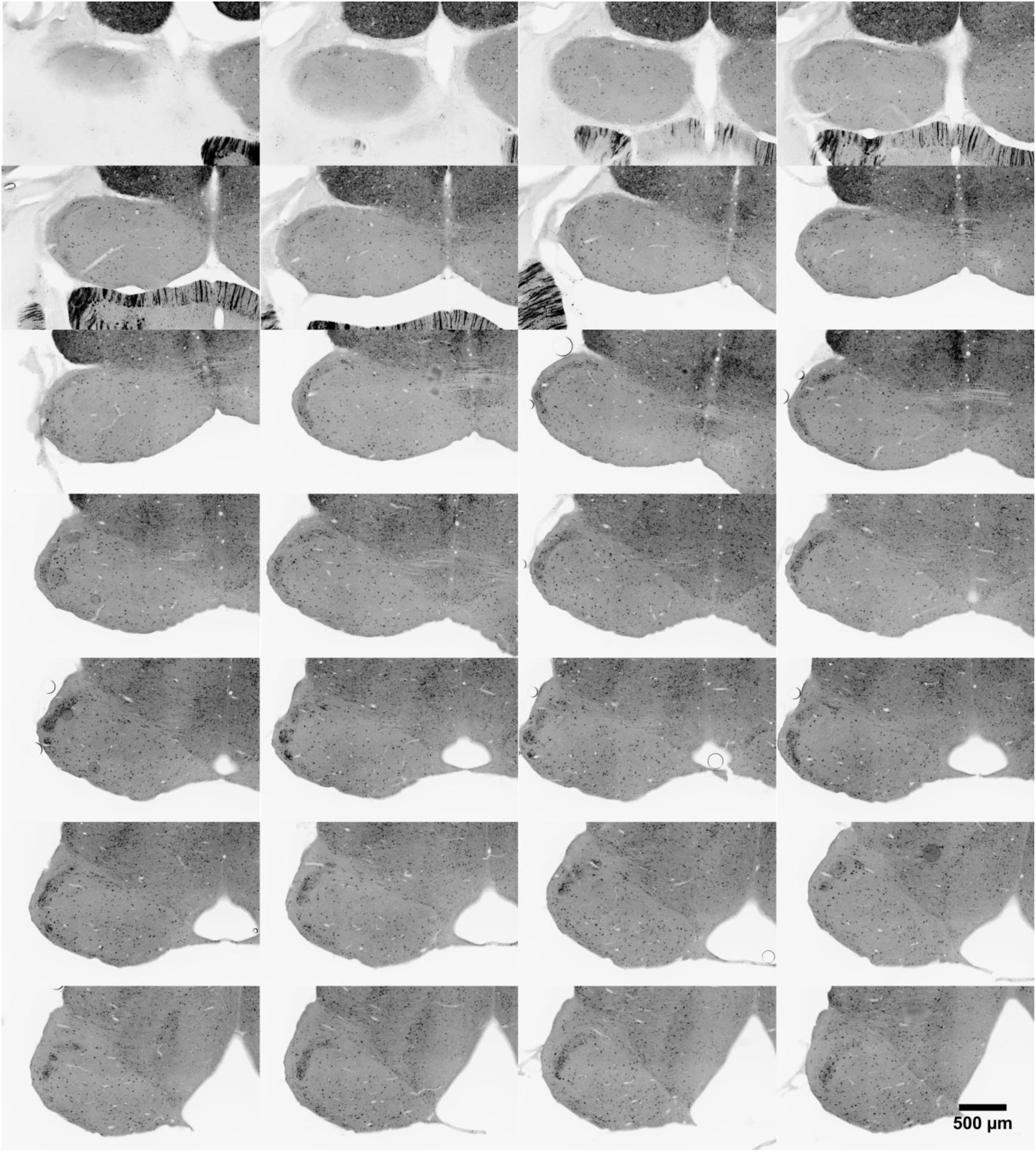
GFP staining of GCaMP6s in the inferior colliculus for the series of consecutive 40 μm horizontal brain slices from the same animal in Figure 9A, displayed from dorsal to ventral (left to right, top to bottom). Same sections as supplement 1.

Video 1

Video of all sound-responsive regions-of-interest plotted in a common coordinate system, color coded according to their characteristic frequencies. The Z-dimension indicates the depth from pia surface. Numbers on the axes are in μm. The same data appears in Figure 6A.

Video 2

Video showing the 3D reconstruction of the most dorsal aspect of the left and right inferior colliculi in animal 12156-04 from 40 μm serial sections. GAD67-dense modules are marked in both ICs. The same sections used for reconstruction are shown in Figure 9A and Figure 9 – figure supplements 1-3.

